# Human outperform mouse Purkinje cells in dendritic complexity and computational capacity

**DOI:** 10.1101/2023.03.08.531672

**Authors:** Stefano Masoli, Diana Sanchez-Ponce, Nora Vrieler, Karin Abu-Haya, Vitaly Lerner, Tal Shahar, Hermina Nedelescu, Martina Francesca Rizza, Ruth Benavides-Piccione, Javier DeFelipe, Yosef Yarom, Alberto Munoz, Egidio D’Angelo

**Affiliations:** Department of Brain and Behavioral Sciences, University of Pavia, Italy; Brain Connectivity Center, IRCCS Mondino Foundation, Pavia, Italy; Centro de Tecnología Biomédica (CTB), Universidad Politécnica de Madrid, Spain; Departamento de Biología Celular, Universidad Complutense de Madrid, Spain; Department of Neurobiology and ELSC, Edmond J. Safra Campus, The Hebrew University of Jerusalem, Israel; Scripps Research Institute, La Jolla, USA; Instituto Cajal (CSIC), Spain; Department of Neurosurgery, Shaare Zedek Medical Center

## Abstract

Purkinje cells (PC) of the cerebellum are amongst the largest neurons of the brain and have been extensively investigated in rodents. However, their morphological and physiological properties in humans are still poorly understood. Here, we have taken advantage of high-resolution morphological reconstructions and of unique electrophysiological recordings of human PCs *ex vivo* to generate computational models and estimate computational capacity. An inter-species comparison showed that human PCs had similar fractal structure but were bigger than mouse PCs. Consequently, given a similar spine density (2/μm), human PCs hosted about 5 times more dendritic spines. Moreover, human had higher dendritic complexity than mouse PCs and usually emitted 2-3 main dendritic trunks instead than 1. Intrinsic electroresponsiveness was similar in the two species but model simulations revealed that the dendrites generated ~6.5 times (n=51 vs. n=8) more combinations of independent input patterns in human than mouse PCs leading to an exponential 2^n^ increase in Shannon information. Thus, while during evolution human PCs maintained similar patterns of spike discharge as in rodents, they developed more complex dendrites enhancing computational capacity up to the limit of 10 billion times.

## INTRODUCTION

Current knowledge on neuronal functions still almost entirely relies on rodents ^1–3^ in which refined electrophysiological and imaging recordings *ex vivo* are routinely used to determine membrane potential changes, dendritic processing, synaptic transmission, and long-term synaptic plasticity. Among the few cases addressing the functional properties of human neurons, studies on pyramidal cells have shown higher dendritic compartmentalization, faster dendritic integration and stronger dendritic amplification in humans compared to rodents ^4–7^. However, no studies have been reported so far about neurons of the human cerebellar cortex.

PCs were first described by Johan Evangelista Purkinje in 1837 ^8^ and then in more detail by Camillo Golgi in 1882 and Santiago Ramón y Cajal, who in 1888 discovered that PC dendrites have dendritic spines ^9^. The PC is among the largest and most complex neurons of the brain and shows typical morphological and electrophysiological properties ^10–22^. In rodents, PCs express a rich complement of ionic channels, synaptic receptors, and intracellular transduction mechanisms that are differentially distributed over the neuron subdivisions. The dendritic tree is almost planar and exhibits a series of ramifications, collecting synaptic inputs and conveying currents to the action potential (AP) initiating site in the axonal initial segment (AIS). Rodents PC dendrites are reported to receive in the order of 10^4^ excitatory synapses from parallel fibers (pf) and 10^3^ inhibitory synapses from stellate cells (SC) ^23,24^, and are thought to operate as a perceptron ^25^ through a process of linear encoding ^26^. For rodents, advanced computational models have been constructed and used to simulate neuronal responses both in isolation ^25,27–36^ and inside microcircuit reconstructions ^24,37^. Little is known about human PCs, which look bigger than rodent PCs ^38^ but do not have any electrophysiological recordings available. So, the first questions to answer are how human PCs discharge and how their structure impact on computational capacity ^39–41^.

As human datasets are far from complete, computational models can be used to fill the gaps caused by missing knowledge and propose specific functional hypotheses ^42^. Detailed neuron models can be reconstructed from digital morphologies to generate morpho-electrical equivalents ^43,44^, which can be subsequently endowed with cell-specific ionic conductances and simulated ^45^. Simulations of responses to current injection can then be optimized against electrophysiological recording templates to extract the missing information about model free parameters, i.e., maximum ionic conductances ^46^. In this work, we have taken advantage of high-resolution morphological reconstructions of human PCs and of unique electrophysiological recordings obtained in acute cerebellar slices from post-surgical cerebellar specimens to generate detailed biophysical models. These allowed us to simulate the electrophysiological response of human PCs in conditions that would otherwise be impractical to assess experimentally and to evaluate their dendritic complexity and computational capacity.

## METHODS

Morphological reconstructions and electrophysiological recordings were performed in both human and mouse PCs and simulations were performed using multicompartmental models (see Figs 1-4).

### Human experimental data

#### Patch clamp recordings

Ex-vivo human cerebellar cortical tissues were obtained from surgeries aimed at deeper brain structures, following the guidelines of the institutional ethics committees and the Helsinki committee of Shaare Zedek hospital, which also granted approval. Tissue samples were considered neurologically normal, as confirmed by a pathologist present during the surgery. Following excision from the brain, cerebellar cortical tissue was placed in ice-cold oxygenated ACSF composed of (in mM) NaCl 126, KCl 2.5, MgSO4 1.5, KH2PO4 1, NaHCO3 24, glucose 10 and CaCl2 1, and bubbled with carbogen (95%O2/5%CO2) to maintain oxygenation and pH. Following transportation to the laboratory, the pia was removed from the cortical surface as much as possible (but without tearing the tissue apart) and tissue chunks were trimmed and/or cut where necessary to expose as much as possible of the translobular plane, while the tissue was maintained submerged in ice-cold oxygenated ACSF. The tissue was then dried briefly and fixed to the stage of a Leica VT1200S vibratome using superglue alongside a piece of agar for structural support. The tissue was then quickly submerged again in ice-cold ACSF continuously bubbled with carbogen, and slices were cut at 300 - 400 μm thickness at a speed of 0.01 - 0.04mm/s and amplitude of 1.25mm. Slices were incubated for at least 1 hour at room temperature in the same solution, and then placed in a warmed (~32degrees C) recording chamber superfused with oxygenated recording ACSF (same solution as for slicing except with 2.4mM CaCl2). Slices were visualized under infrared light in an upright microscope with 40x water-immersion objective, and whole-cell patch-clamp recordings of PCs were established using electrodes with 2 - 6 MΩ resistance pulled from borosilicate glass and filled with an intracellular solution composed of (in mM) K-gluconate 140, HEPES 10, EGTA 0.01, CaCl2 0.001, MgATP 4 and 1% biocytin (w/v), pH adjusted to 7.2 - 7.3 using KOH and osmolarity 290 - 310 mOsm. Electrical activity was recorded in current-clamp mode at a sampling rate of 20-50kHz and low-pass filtered at 10kHz using a MultiClamp 700B amplifier and Digidata 1550B digitizer connected to pClamp software, and stored for offline analysis. Images of the recorded neuron’s morphologies were obtained by staining for biocytin using streptavidin conjugated to Alexa-647 and imaging the mounted samples under 10x magnification (see Fig. 1).

**Figure 1.**
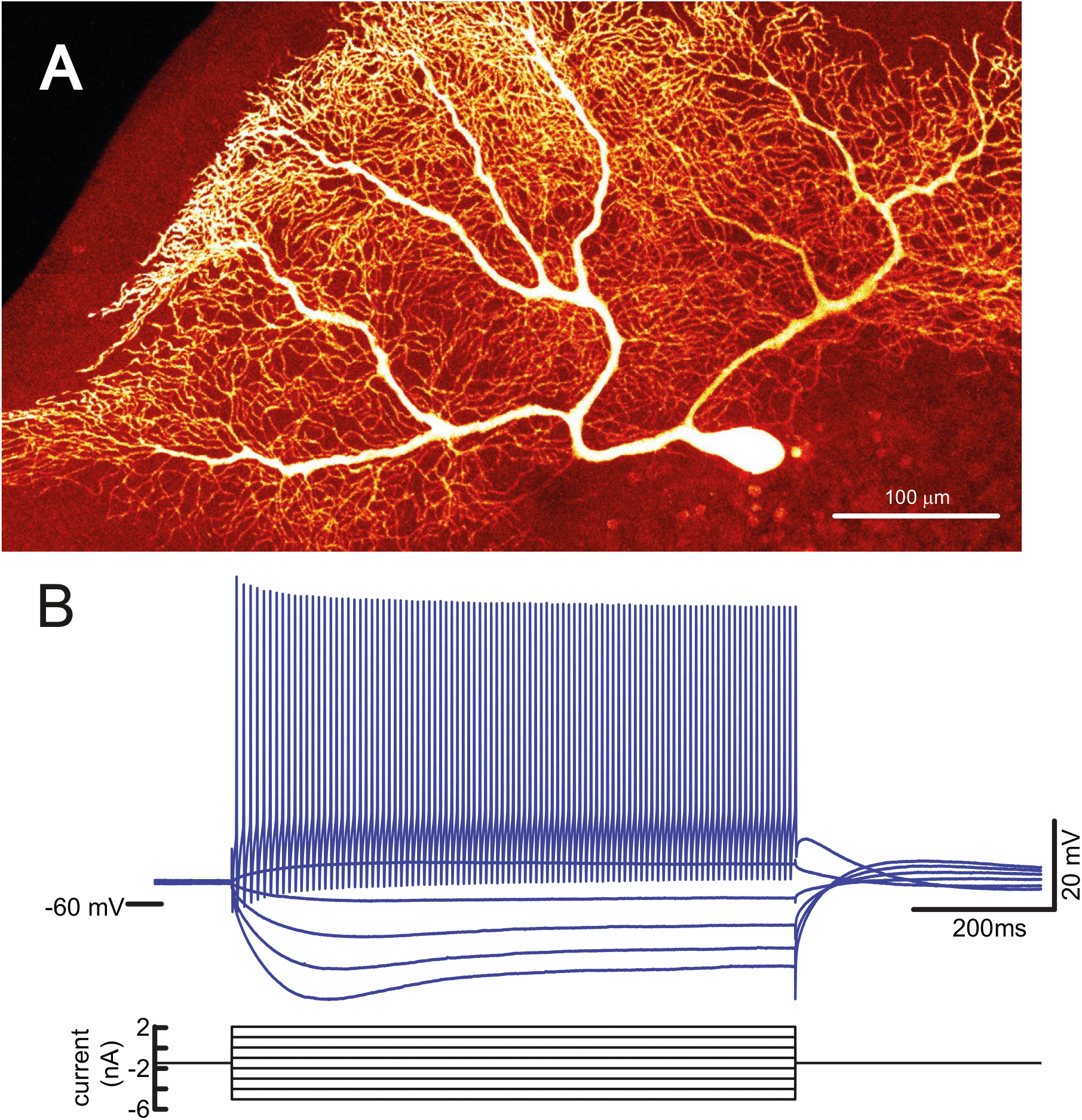
Human PC experimental recordings. (A) Biocytin/streptavidin fluorescence confocal image of a human PC dye-filled with Alexa-647 during a whole-cell patch-clamp recording in an acute cerebellar slice *ex vivo*. Note the characteristic palisade formed by the dendritic tree that stems with multiple trunks from the soma. The dendritic tree extends for almost 1 millimeter on the parasagittal plane. (B) Example of voltage responses to the injection of positive and negative current steps in the same cell shown in A. Note regular firing in the depolarizing direction and sagging inward rectification in the hyperpolarizing direction.

#### Morphological reconstructions

Human brain tissue from two autopsy cases (AB6, a 92 years-old female dead from a heart failure; and AB7, a 66 male dead from a bladder carcinoma) was collected from the Unidad Asociada Neuromax—Laboratorio de Neuroanatomía Humana, Facultad de Medicina, Universidad de Castilla-La Mancha, Albacete (Spain). Brain samples were obtained following the guidelines of the Institutional Ethical Committees, which also granted approval. Cerebellar tissue was considered normal since AB6 and AB7 individuals were free of neurological and psychiatric diseases, although abundant AT8 positive cells were found in the cerebral cortex of the AB6 case ^47^. The postmortem delay between death and tissue processing was below 4 h (AB6) and 2,7 h (AB7). Upon removal, the brains were immediately fixed in cold 4% paraformaldehyde in phosphate buffer (PB 0.1 M, pH7.4) for 24 h. Small blocks from the cerebellar vermis were cut from which parasagittal sections (300 μm) were obtained with the aid of a Vibratome (Leica).

For intracellular injections, sections were prelabeled with 4,6-diamidino-2-phenylindole (DAPI; Sigma, St Louis, MO), and a continuous current was used to inject with Lucifer yellow (8 % in 0.1; Tris buffer, pH 7.4; LY) individual PCs from the vermis of the anterior and posterior lobes. LY was applied to each injected neuron by continuous current until the distal tips of their branches fluoresced brightly, indicating that the dendrites were completely filled and ensuring that the fluorescence did not diminish at a distance from the soma. Following the intracellular injections, sections were incubated for 72h at 4°C in stock solution (2% bovine serum albumin, 1% Triton X-100, and 5% sucrose in PB) containing rabbit anti-LY antibody (1:400 000; generated at the Cajal Institute, CSIC, Madrid, Spain). The sections were then rinsed in PB and incubated in biotinylated donkey anti-rabbit IgG (1:100; Amersham, Buckinghamshire, United Kingdom). Then, sections were rinsed again and incubated Alexa fluor 488 streptavidin-conjugated (1:1000; Molecular Probes, Eugene, OR, United States of America). Finally, the sections were washed and mounted with ProLong Gold Antifade Reagent (Invitrogen Corporation, Carlsbad, CA, USA). See ^48,49^ for further details of cell injection methodology.

For cell reconstruction and quantitative analysis, sections were imaged with a confocal scanning laser attached to a fluorescence microscope Zeiss (LSM710). Fluorescent labeled profiles with different wavelengths were recorded though separate channels. Stacks of images at high magnification (×63 glycerol; voxel size, 0.057 × 0.057 × 0.14 μm^3^) were acquired to capture dendritic arbors on the basis of LY immunostaining. Since intracellular injections of PCs were made in 300 μm-thick parasagittal sections, the part of the dendritic arbor nearest the surface of the slice from which the cell soma was injected (typically at a depth of ~30-50 μm from the surface) could be partially lost. In our case, most of the dendritic arbor is estimated to be included within the section since we only reconstructed PCs with dendritic trees running parallel with respect to the surface of the parasagittal slice, and the geometry of dendritic trees of PCs is largely flat and restricted to a parasagittal plane ^50^ (see Fig. 1 Supplementary).

Data points of neuron morphology of each PC included in the analysis (n=6) were extracted in 3D using Neurolucida 360 (MicroBrightfield). Briefly, dendrites, axon and soma, in the skeleton definition were described through 3D points, delimiting the different segments that form the cell arbor. These points have an associated diameter that provides the information of the varying thickness of the dendritic or axonal processes at that particular point, and varies along the length of the processes. The soma was defined through a set of connected points tracing the contour of the soma in 2D.

To calculate the number and density of spines, spines from all sizes and morphologies were counted in dendritic segments with diameters decreasing in intervals of 0.1 μm (ranging from 1,2 to 0,2 μm) at different points along the length of dendrites from 5 reconstructed PCs. Spine density values in dendrites with different diameters were obtained by dividing spine counts per the length of each segment. These measured density values were used to estimate the total number of spines and the mean density value in PCs, taking in consideration the length of dendritic branches with different diameters over the entire dendritic tree. Dendritic spine structure was analyzed using Imaris 6.4.0 (Bitplane AG, Zurich, Switzerland) in a selection of 150 dendritic spines. Since confocal stacks of images intrinsically result in a z-dimension distension, a correction factor of 0.84 was applied to that dimension. This factor was calculated using a 4.2μm Tetraspeck Fluorescent microsphere (Molecular Probes) under the same parameters used for the acquisition of dendritic stacks. No optical deconvolution was used for spine reconstruction. Spine head area was 3D reconstructed in a selection of spines showing a clear head, whose morphology could be captured using a single surface of a particular intensity threshold. The spine neck length and spine neck diameter were manually marked in each selected dendritic spine, from the point of insertion in the dendritic shaft to the spine head, while rotating the image in 3D. (see ^49^ and ^51^ for further details) (see Fig. 2).

**Figure 2.**
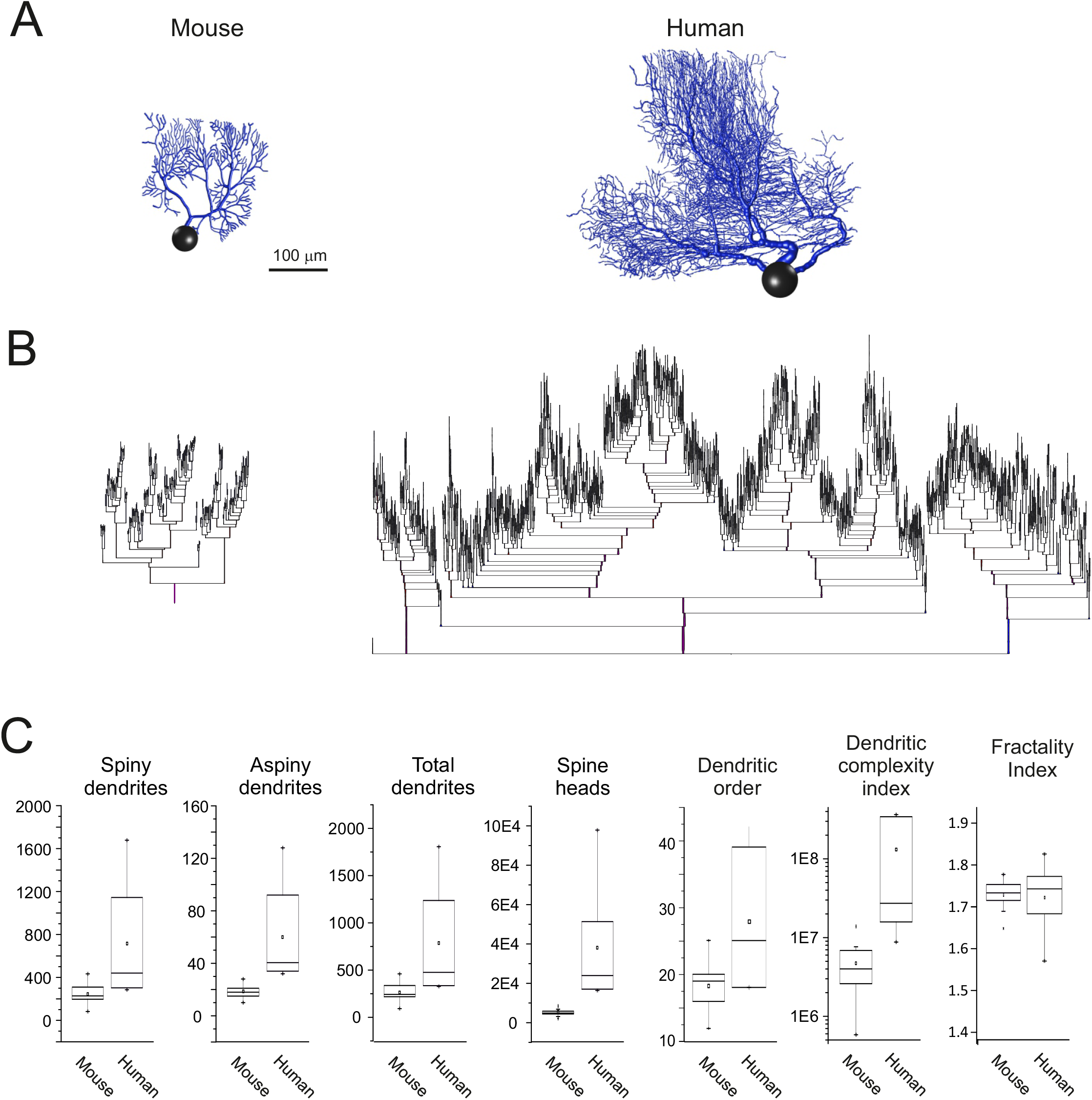
Human and mouse PC morphological properties. (A) Morphological reconstructions of a mouse PCs and a human PCs derived from fluorescent confocal images of cells injected from postmortem fixated brains and injected with Lucifer yellow (see Fig 3 Supplementary for more reconstructions). Note the similar shape but greater size of human than mouse PCs and the characteristic shape of the dendrite. (B) Dendrograms of a mouse and human PC. The PCs dendrograms showed similarities in the architecture of dendritic ramification. The insets magnify two terminal regions of dendrograms highlighting the organization of spiny dendrites (<3.5μm in diameters), which account for approximately 90% of segments. All the comparisons between human and mouse PC parameters reported in the figure reveal a statistically significant difference at p<0.01 (unpaired *t*-test), except for the fractality index. (C) The boxplots show the metrics of reconstructed dendritic morphologies (19 mice and 6 human PCs). These include the number of spiny and aspiny dendrites, total dendrites, spike heads, and dendritic order. Note the larger numbers for all parameters in human than mouse PCs (all differences are statistically significant, see text). The dendritic complexity index (DCI) shows about one order higher complexity in human than mouse PCs. The fractality index shows similar values in human and mouse PCs (colored dots represent individual values for all reconstructed PCs).

### Mouse experimental data

#### Patch-clamp recordings

Mouse recordings were carried out on 26-30 day-old (P0=day of birth) C57BL/6N wild-type mice cerebellar slices. The mice were anaesthetized with halothane (Sigma-Aldrich) and killed by decapitation to remove the cerebellum for acute slice preparation according to a well-established technique. The cerebellar vermis was isolated and fixed on the vibroslicer’s stage (Leica VT1200S; Leica Biosystems) with cyano-acrylic glue. Acute 220 μm-thick slices were cut in the sagittal plane and, during the slicing procedure, the cerebellar vermis was immersed in a cold (2-3 °C) oxygenated bicarbonate-buffered saline solution (Kreb’s solution) containing (mM): NaCl 120, KCl 2, MgSO4 1.2, NaHCO3 26, KH2PO4 1.2, CaCl2 2, glucose 11 (pH 7.4 when equilibrated with 95%O2-5%CO2). Slices were incubated at room temperature in oxygenated standard extracellular solution for at least 1 h until use. For whole cell patch clamp recordings, slices were placed in a chamber continuously perfused at a rate of 1.5 ml/min with oxygenated Kreb’s solution and maintained at 32°C with a Peltier feedback device (TC-324B, Warner Instrument Corp.). Slices were visualized using an upright epifluorescence microscope (Axioskop 2 FS; Carl Zeiss) equipped with a 63, 0.9 NA water-immersion objective (Olympus, Hamburg, Germany). Whole-cell patch-clamp recordings from the soma of PCs were performed with Multiclamp 700B [-3dB; cutoff frequency (fc), 10 kHz], sampled with Digidata 1550 interface, and analyzed off-line with pClamp10 software (Molecular Devices). Patch pipettes were pulled from borosilicate glass capillaries (Sutter Instruments) and filled with internal solution containing (in mM): potassium gluconate 126, NaCl 4, HEPES 5, glucose 15, MgSO4 7, H2O 1, BAPTA-free 0.1, BAPTA-Ca2+ 0.05, Mg2+ 140 -ATP 3, Na+ -GTP 0.1, pH 7.2 adjusted with KOH. Pipettes had a resistance of 2–3 MΩ when immersed in the bath. Signals were low-pass filtered at 10 kHz and acquired at 50 kHz.

All procedures were conducted by European guidelines for the care and use of laboratory animals (Council Directive 2010/63/EU) and approved by the ethical committee of the Italian Ministry of Health (628/2017-PR).

#### Morphological reconstructions

The 3D morphologies of 19 PCs were reconstructed from the anterior zone of lobule V of post-natal (P)27 L7-tau-GFP mice using Neurolucida (MBF Bioscience) ^22,52^. To control for inter-animal variation in anatomy such as folia size, age-matched male mice were used. Briefly, following transcardial perfusion (4% paraformaldehyde in 0.1 M phosphate buffer), brains were removed from the skulls, postfixed overnight in the same fixative and transferred to a 0.01 M phosphate buffer saline (0.9% sodium chloride) solution (PBS). A Leica vibratome was used to collect 80 μm thick parasagittal sections in cold 0.01 M PBS. Subsequently, tissue sections containing lobule V near the midline in the medial-lateral plane were mounted onto glass slides, air dried and cover-slipped with antifade Prolong mounting medium (Invitrogen). An LSM 710 Zeiss confocal microscope equipped with a 63X oil objective lens (NA 1.46) was used to collect confocal stacks using ZEN imaging software. In order to unambiguously resolve dendrites of neighboring PCs in the parasagittal plane, the following image acquisition parameters were used: *x, y*-size of 0.22 *μ*m/pixel; dimensions *x:* 224.7 *μ*m, *y:* 224.7 *μ*m; image size 1024 × 1024) and a *z*-step size of 0.25 *μ*m.

Confocal image stacks were imported into the Neurolucida software connected to a Wacom digital tablet and pen. Dendritic tree reconstructions were generated using the interactive and manual tracing functions, which inserted points along each dendritic branch. The location of these points and their coordinates in 3D space (x, y and z) thus described the geometric shape of each dendritic arbor. Somata were drawn with Neurolucida’s continuous tracing function, which inserted points along the contours of somata as visualized in each plane of section (Fig. 2 Supplementary).

### Modelling

The modelling workflow was developed in Python3 and NEURON8 ^45^ and the subsequent optimization was performed using the “Blue Brain Python Optimization Library” (BluePyOpt) ^46^. Here we report an advanced workflow for modelling and simulation of human and mouse PCs. All models were optimized at 32° to match the experimental data (see Fig. 3A).

**Figure 3.**
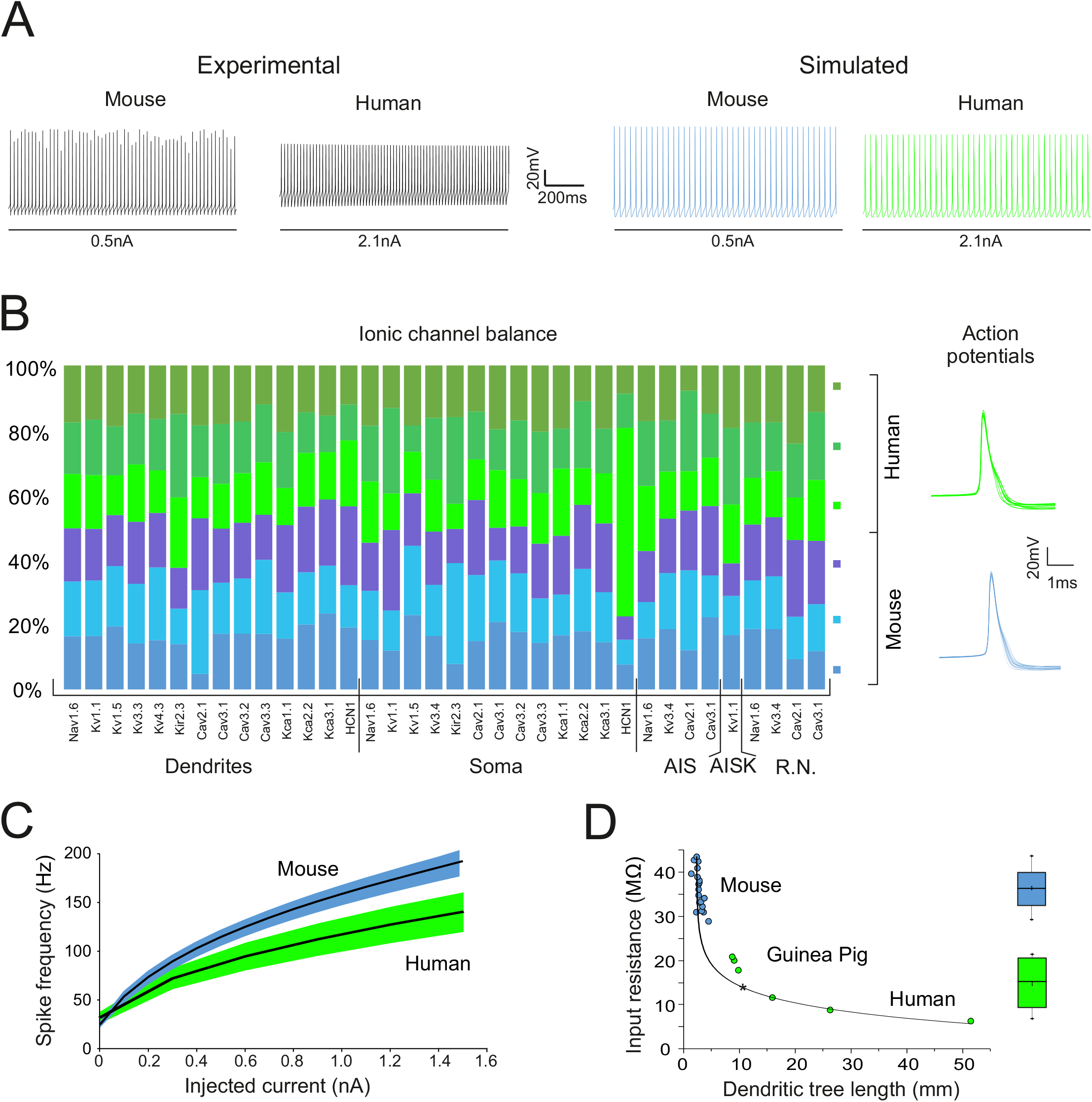
Electrophysiological recordings and model simulations. (A) Left: experimental current-clamp recordings from a mouse and a human PC injected with positive current. The two cells generated similar responses, but the human PC required a higher current injection. Right: simulations in a mouse and a human PC model injected with the same positive and negative current steps used in the experiments. The mouse and human PC models generated similar responses, which closely corresponded to those recorded experimentally. The higher current injection required by human than mouse PC is due to the correspondingly higher input conductance. Trace colors correspond to the cells reported in panel B. (B) The bar plot shows the balance of maximum ionic conductances in 6 randomly chosen models, 3 from mouse and 3 from human PCs. For each conductance, the bar colors show the % ratio among cells. Note that the balance is quite similar for mouse and human PCs for most ionic channels (one noticeable deviation is visible for HCN1 in one of the cells). The traces on the right show action potentials in the mouse and human PC models. Except for minor deviations in the AHP trajectory, the action potentials were almost identical. (C) Average I-F relationships for mouse and human PC models. The I-F slope was 2.05 ± 0.14 times higher in mouse than human models reflecting higher input resistance (see panel E). The shadowed areas represent s.d.. (D) Dependence of input resistance on dendritic length (input resistance was estimated in all the PC models using voltage-clamp step protocols). Input resistance is inversely correlated with the dendritic tree length along an exponential decaying function, i.e., PCs with longer dendrites have smaller input resistance. Human PCs lay in the right-hand branch of the curve, mouse PCs in the left-hand branch, the guinea-pig PC ^28,36^ is in the middle. The box plot on the right shows statistics of input resistance for all the mouse and human PC models.

#### Electrotonic structure and multi-compartmental neuron organization

Morpho-electric equivalents (Fig. 2) were generated from the mouse and human PC morphologies obtained from Neurolucida. The mouse morphologies required curation to reduce unevenness in the main dendritic trunks, while the human morphologies were used without any modifications (for all the morphologies see Fig. 3 Supplementary, Table 1 Supplementary). The soma dimensions were calculated using the largest contour of each cell and substituted with an equivalent spherical shape. During model optimization, both mouse and human PC morphologies were attached with the same axonal compartments, composed of an AIS and a myelinated axon, containing 3 Ranvier’s nodes ^28,36^. The dimensions of the human PCs AIS was kept the same as in mice because, in two out of six reconstructions, the average length and diameters were in line with mouse data ^53^. For consistency with previous models ^36^, the axon collateral was also added.

The dendrites of previous PC models ^36^ were subdivided into spiny and aspiny based on the dendritic properties of Guinea Pigs ^30^. Here we adopted the same criteria based on the following parameters: dendritic length, dendritic diameter, total number of dendrites, total length of the dendritic trees. Based on the conserved shapes and morphological properties, as shown in different animal types ^54^, the number of segments belonging to each zone was carefully rescaled. The following distribution allowed to maintain the same proportion of spiny and aspiny dendrites throughout all PC morphologies in mice and human as well as in Guinea pig PCs: the segments having a diameter below 1.6μm were labelled as spiny dendrites (distal dendrites), the segments between 1.6μm and 3.3μm were labelled as part of the trunk (proximal dendrites), and the segments above 3.3μm were labelled as part of the main trunks containing sodium channels.

#### Passive properties

The passive properties of mouse and human PC models were derived from those of the Guinea Pig model ^28,36^. The axial resistance was kept at Ra=122 Ωcm. The membrane capacitance was set to C_m_= 1μF/cm^2^ for the soma and C_m_= 2μF/cm^2^ for dendrites larger than 1.6μm. In agreement with previous models ^27,29,36^, the compartments below 1.6 μm received a variable C_m_ value to compensate for the spines when these were omitted from the models according to the following equation:

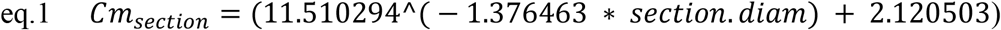

In a first iteration of this workflow, C_m_ was set as a free parameter ^55^ but, to reduce the parameter space, it was later fixed to the value given by eq.1 ^56^. The leak values were set to 0.003 S/cm^2^ for the soma and to 0.0003 S/cm^2^ for the remaining compartments, except for the myelinated compartments, which received the NEURON passive channel (pas). The leak reversal potential was set, for all compartments and morphologies, to −61mV.

#### Active membrane properties

The distribution of the ionic channels, their reversal potential, the calcium buffer and pump densities were taken from previous PC models ^28,36^. This choice proved effective but minor adjustments were required to comply with recent experimental data on mouse and human PCs. The Kv1.1/2 potassium channels were previously distributed on the somato-dendritic compartments, based on rat experimental data ^57^, while the latest immunohistochemical data indicated that channel distribution was limited to the main dendritic trunk ^58,59^. The latest experimental data about Kv1.5 showed a somato, proximal dendritic distribution ^58,59^ compared to previous report with the channel on the entire dendritic tree. The Kca2.2 potassium calcium-dependent channel was previously confined to a part of the dendritic tree, whereas in mice can be found on the entire tree ^60^. The calcium buffer was improved with the calcium protein Calmodulin ^61^ which was previously validated in two other cerebellar models ^56,62^. The distribution of ionic channels among compartments, including spines, is shown in supplemental Fig. 4 Supplementary.

#### Synaptic mechanisms

Based on previous PC models ^28,33,56^, three synaptic zones were defined. 1) Dendrites from 0 to 0.75μm were targeted by ascending axons, 2), dendrites from 0.75μm to 1.6μm were targeted by pf and 3) dendrites from 1.6μm were targeted by climbing fibers (cf). The GABAergic synapses were distributed on compartments from 0.3 to 1.6μm ^63^. The membrane mechanisms for both AMPA and GABA receptors were the same as in ^28^ with a peak synaptic conductance of 2600pS each. The synapses contain a presynaptic vesicle cycle and neurotransmitter release mechanism, a 2D diffusion process and Markov chain molecular models to simulated the postsynaptic receptor ^64^.

#### Feature extraction from experimental data

The data were extracted from mouse and human PC experimental traces having spontaneous activity and responses to positive step current injections (mouse PCs: 0.1, 0.5 and 1nA; Human PCs: 0.1, 0.5 and 1nA). The Blue Brain Project eFel library ^65^ was used to extract the following feature: AP amplitude, ISI_CV, AHP depth slow, AHP depth absolute, AP width, voltage base, mean frequency and Spike count.

#### Optimization of maximum ionic conductances

Maximum conductance parameters optimization was performed with the open source “Blue Brain Python Optimization Library” (BluePyOpt) ^46^ (Fig. 5 Supplementary, Table 2 Supplementary, Table 3 Supplementary, Table 4 Supplementary). The initial population was set to 288 individuals, which yielded a final population of 576 individuals. The optimizations were initially performed for a dozen of generations and repeated to find the best solutions. The final optimizations, which generate the data used for the validation process, lasted for only 7 generations and reached a near zero fitness value with just 3-4 generations. Optimizations were run on the Piz Daint cluster (CSCS - Lugano), using 8 nodes with 36 cores each, for a total of 288 cores. Simulations were performed using variable time-step, model optimizations with fixed time step (0.025 ms). The smallest morphology (a mouse PC morphology with 103 compartments), was optimized in 2 hrs, whereas the biggest one (a human PC morphology with 1880 compartments) was optimized, on average, in 19hrs.

#### Model simulations of optimized models

The final populations obtained from optimization were simulated using the BluePyOpt template enriched with a custom-built MPI code (Mpi4py), to exploit all available CPU cores. This allowed the distribution of the simulations across 8 nodes (288 cores) with a total simulation time ranging from 22 min to 3 hrs to simulate the last generation of each optimization.

#### Validation workflow

The validation was performed on Piz Daint (CSCS - Lugano) using a variable number of nodes, depending on the number of validated individuals. The workflow was subdivided as follows:

1. The simulated voltage traces were analyzed to assess the presence of spontaneous firing activity (0nA) and the I/O relationship (0.5 and 1nA). Based on the experimental data, a valid model should have spontaneous firing with a frequency between 5Hz and 50Hz, in compliance with Zebrin plus PCs ^66,67^. To complete the I/O relationship, the maximal frequency for the 0.5nA and 1nA currents was set to 150Hz and 200Hz, respectively.
2. Sodium channels in the AIS are critical for AP generation ^36,68^. The models were simulated for 1s after removing sodium channels in the AIS. The models that, in this configuration, lost their ability to generate APs were validated and passed to the next phase.
3. The Input resistance (Rin) was assessed using the NEURON Voltage Clamp MOD file. Since it was not included in BluePyOpt, it was added in the template and configured to record the current for each simulation. The models were simulated for 300ms, with a square current (−70mV, −80mV, −70mV) with each step lasting for 100ms. Rin was calculated automatically, and models were validated when Rin fell in the experimental range.
4. The last validation step required a series of modifications to the template to accommodate AMPA receptors, and new compartment lists to use synaptic inputs instead of injected currents. Each model was provided with 50 excitatory synapses, one for each compartment belonging to the pf compartment list. We examined four bursts frequency: 50, 100, 200 Hz, each one composed of 10 spikes. The bursts were delivered 5 times, each time with a delay of 1s, for a total simulation time of 5.5s. Features were extracted from the last burst, using a mobile time window to include only the spikes generated by the synaptic bursts. The resulting voltage and current traces were stored in HDF5 files. A model was validated with 9-10 spikes at 50Hz, 4-6 spikes at 100Hz, 3-4 at 200Hz (Table 2 Supplementary).

Validated models were randomly chosen and imported into a custom-made Python3/NEURON 8 script. Python allowed a higher degree of freedom in checking synaptic patterns, automatic currents recording, graph creations, etc. The simulations were carried out with Python3, NEURON 8, at 32°, with a fixed time step (0.025), on an AMD 1950x 16core/32threads.

#### Spine morphology and simulations

To analyze the impact of localized synaptic patterns, spines were added to the PC models. Each spine was constructed with two cylindrical compartments to represent the head and the neck. The dimensions of the head were 0.35μm in length and 1μm wide, whereas the neck was 0.7μm long and 0.2μm wide ^69,70^. The measured human values were similar, with a head length of 0.26 and 1μm wide, whereas the neck was 0.72μm long and 0.18μm wide. The neck C_m_ was set to 3μF/cm^2^ instead, the head CM was set to 1μF/cm^2^. The ionic channels were placed only on the spine heads: Cav2.1 (P-type), Kca1.1 (BK) Kca2.2 (SK2) ^60,71^, Kv4.3 (A-type), Cav3.1 (T-type) ^72,73^. The calcium channel Cav2.3 was experimentally proven not to be critical for intrinsic and synaptic responses, so it was omitted ^72^. Given the importance of calcium dynamics, the Ca-buffer was inserted in the spines too. The average experimental distribution indicated 2 spines/μm ^74–76^ and the same was used for humans (internal experimental data). In this model configuration, C_m_ for the entire dendritic tree was set to 2μF/cm^2^ regardless of compartment diameter ^27–29,36^.

#### Spines synaptic stimulation

To activate a specific synaptic cluster, we developed a custom Python script automatically dividing each morphology into a square grid of 30*30μm (900μm^2^), containing spines belonging to aa, pf and SC. The simulations were performed with the same synaptic patterns previously described and applied to a randomly chosen 30% of all the available spines ^77^. The computational impact of the extended model required the usage of a single blade cluster, composed of two AMD 7501 processors containing 32 cores each. The memory impact, even during the longest simulation, was about 10-15GB. The mice simulations were better suited to run on 16 cores, whereas the human required 32 cores. The temperature and time step remained the same as previously defined.

### Data analysis

Part of the results were analyzed with custom Python3 scripts and, for specific cases, with MATLAB 2018b ^78^ provided by the University of Pavia. The dendrograms were generated with NeuroM (BlueBrain Project (BBP) and the morphological properties were analyzed with Matlab-based module, “the TREES toolbox” ^79^.

The fractal index was calculated with the Sierpinski triangle methods ^80^ with a range from 0.01 to 1 (40 logarithmic scales). To provide the best possible resolution for the triangle methods, the morphologies were plotted with Vaa3D, rescaled accordingly to their somatic dimension, and saved into images of 1024*1024 pixels.

The 3D plots of the morphologies were generated with Vaa3D ^81^. The dendritic complexity index (DCI) ^82,83^ was determined from the following equation:

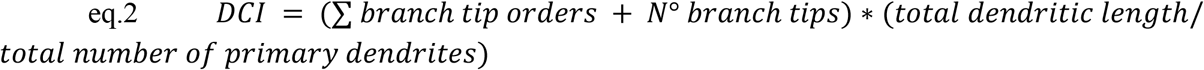

All these measurements are readily available from our automated analysis of dendrograms. Branch tip order is an integer value assigned to each terminal tip of the neuron’s dendrite, equaling the number of sister branches emanating from the dendritic segment between a particular terminal tip and cell body.

Impedance was calculated with NEURON using the built-in “impedance” class. The soma was set as the target and one dendrite every 20 was set as the source of the transfer function. Membrane voltage changes were elicited by a sinusoidal current at 10Hz injected into a dendritic compartment. This allowed to calculate the *input conductance* at the injection site and the *transfer conductance* at other dendritic sites ^84^.

### Model dissemination

The mouse morphologies will be uploaded to neuromorpho.org and the computational models will be made available on eBRAINS and ModelDB. The human morphologies and computational models will be made open source.

## RESULTS

### Human and mouse PC morphological properties

Since early on, rodent PCs have been intensely investigated ^10,11,15,17,19,20,25,27–32,34,36^ whereas the properties of human PCs have remained undisclosed until now. Fig.1 shows the first recordings from human PCs in cerebellar slices obtained from fresh postsurgical specimens. The human PCs showed morphologies (Fig. 1A) and electroresponsive patterns (Fig. 1B) similar to those observed in rodents, although some striking differences were the greater dendritic tree and the higher current needed to elicit action potentials in somatic current-clamp recordings.

To get insight into these properties, we carried out a detailed comparison of human and mouse PCs aided by computational models. We considered 19 mouse and 6 human PC 3D morphologies reconstructed using fluorescence confocal microscopy (Fig. 2A) and evaluated several metrics on dendrograms to unveil the architecture, hierarchies and dependencies of dendritic branching (Fig. 2B; Table 1 Supplementary). The dendrograms recapitulate the dendritic structure in a graph made of segments stemming from branching points. In both species, above 90% of all segments belonged to spiny dendrites. The area covered by PCs was 5.5 times larger in human than mouse PCs (Fig. 2A), with a maximum size of 175500 μm^2^ in the former and of 29700 μm^2^ in the latter. Moreover, in mice, 17/19 PCs had a single dendritic trunk, while the remaining 2/19 PCs had two dendritic trunks stemming from the soma ^21,22^. In humans, 1/5 PCs had a single dendritic trunk, 2/5 had 2 dendritic trunks and 2/5 PCs had 3 dendritic trunks stemming from the soma. Thus, human had a much higher probability than mouse PCs (80% vs. 10.5%) to have multiple dendrites stemming from the soma (see also Fig. 2 Supplementary).

The average length of the dendritic tree was 2782 ± 7 μm (n=19) in mice and 20166 ± 2 μm (n=6) in human PCs, so that human were 7.2 times longer than mouse PC dendrites. In comparison, PCs of the guinea pig ^30^, one of the biggest rodents, showed dendrites (9122.6 ± 1224.9 μm; n = 3) 3.27 times longer than mouse PCs ^85,86^. The total number of segments, the number of segments belonging to spiny dendrites, and the number of segments belonging to aspiny dendrites were 2.9, 2.9, and 3.3 times larger in human than mouse PCs, respectively (Fig. 2C). The average spine density was similar in human and mouse PCs (2/μm) but, since the total length of spiny dendrites was higher, the calculated total number of spines was 5 times higher in human than mouse PCs (Fig. 2C) rising from a median of ~5000 to ~25000 spines (ranging up to 100000 in a case). The dendritic branching orders had a broader range in humans (18-41) compared to mice (12-25) (Fig. 2D) reflecting the higher number of intermediate branches needed to demultiplex and scaffold the huge number of terminal branches.

To quantify the impact of branching on PC architecture, we calculated the *dendritic complexity index* (DCI) ^82,83^. This approach, based on purely morphological properties (eq.2 in Methods) demonstrated that human PCs had 6.5 times higher DCI compared to mice PCs (median 26*10^6^ vs. 4*10^6^) (Fig. 2C). Note that DCI in human outperformed that in mouse PCs because of the higher number of sister and terminal branches as well as for the total dendritic length but not for the higher number of primary dendrites (which appears at the denominator in eq.2).

To evaluate the scale invariance of the PC dendrites, we used fractal analysis. The fractal dimension estimated with the Sierpinski triangle method ^80^ was 1.72 ± 0.09 (n=6) in human and 1.73 ± 0.03 (n=19) in mice PCs. The fractal dimension was not statistically different (p=1, unpaired *t*-test) between the two species, suggesting that mouse and humans PC geometries followed a similar scaling rule (Fig. 2C).

### Electrophysiological and biophysical properties of human and mouse PCs

To compare human to mouse PC electrophysiological properties, we used PC current clamp recordings from acute cerebellar slices (Fig. 1 and 3A). A limited number of human PCs were obtained from fresh postsurgical cerebellar tissue preparations. The human PCs (n=3) showed rhythmic activity at rest before breaking into the whole-cell configuration. In current clamp, the PCs were immediately stabilized subthreshold and then injected with depolarizing and hyperpolarizing current steps. In the depolarizing direction, the PCs showed a regular firing pattern that increased in frequency with the injected current, while in the hyperpolarizing direction, the PCs showed sagging inward rectification. Similar properties were observed in mice PCs too. Similar voltage deflections and spike frequency changes were obtained in human and mouse PCs, although the latter required larger current injections. These observations suggested the engagement of similar electroresponsive mechanism both in human and mouse PCs.

### Computational models of human and mouse PCs

Since the information coming from human PC recordings was incomplete, we used computational models to further analyze PC intrinsic electroresponsiveness. Computational models were used to fill the gap in knowledge through principled rules based on neuronal biophysics ^3^ under minimal assumptions.

First, since the human appeared as a rescaled version of the mouse and guinea pig PC (see Figs 1-2), we assumed that the fundamental biophysical properties were also conserved. Thus, we generated both human and mouse PC models starting from the PC model developed for the guinea pig, which represents the gold-standard at present ^27–30,36^.

Secondly, since the spike discharge was similar (see Figs 1 and 3A), the same set of ionic channels was used in human and mouse as well as in the guinea pig PC models (Table 3 Supplementary). This assumption was supported by the expression of equivalent channel genes in these species ^87^. Ionic channels were placed into neuronal compartments of human and mouse PC models as it was done in the guinea pig PC model. The only exceptions were that Kv1.5 was absent from the soma and that Kca2.2 was distributed over the entire dendritic tree in human and mice but not in guinea-pig PC models ^59^.

Importantly, since the electrophysiological traces contain information about the contribution of all ionic channels engaged, the maximum ionic conductances could be parametrized through an automatic optimization process ^46^ based on the comparison between simulated traces and electrophysiological traces used as templates (see Methods). This methodology allowed us to optimally exploit the information contained in human PC recordings. After parameter optimization, the human and mouse PC models did not show relevant differences in spike discharge and ionic conductance balance (a fact that was true also for the guinea pig PC model; ^36^) (table 4 Supplementary) (Fig. 3A-B), reflecting the similar dendritic architecture and ionic channel complement. The spikes generated by these models during spontaneous firing showed near identical shape in human as in mouse PCs (Fig. 3B, right). Likewise, the I-F relationship was also similar but that of mice was steeper (Fig. 3C).

### Distinctive properties of human PC models

I-F relationships were generated for all human and mouse PC models using somatic step-current injections (Fig. 3C; see also Fig. 6 Supplementary). The average I-F was similar, but the human PCs required 2.05 ± 0.14 times more positive current than mouse PC models to reach the same discharge frequencies (e.g., 100 Hz) (Fig. 3D). At 0-current injection, spontaneous firing frequency in mouse PCs (24.2 ± 3.4 Hz; n=19) and human PC models (28.4 ± 8.0 Hz; n=6) was statistically indistinguishable (unpaired t-test, p=0.09).

Input resistance was 36.2 ± 4.3 MΩ (n= 19) in mouse PC models and 14.3 ± 5.6 MΩ (n= 6) in human PC models (Fig. 3D, right). The input resistance value decreased with the total length of the dendritic tree along a continuous exponential distribution independent from species, with the guinea PC model laying in an intermediate position between mouse (the shortest) and human (the longest) PCs (Fig. 3D). The input resistance value of mouse PC models corresponded closely to that reported experimentally ^14^.

### Spontaneous firing and spike backpropagation in PC models

The backpropagation of spontaneously generated APs through the dendritic tree showed remarkable similarities in human and mouse PC models (Fig. 4A). The APs were generated in the AIS, where the AP reached its highest amplitude (40mV). Then, the AP amplitude decreased when the spikes back-propagated to soma and dendrites, reaching just −50mV in the apical dendrites, reproducing a typical PC property ^11^ (Fig. 4B). The AP back-propagation also featured a delay that depended linearly on the distance from soma. Therefore, in the model, the dendritic tree in human and mouse PCs behaved similarly to guinea pig PCs, i.e., it acted as a filter limiting AP back-propagation.

**Figure 4.**
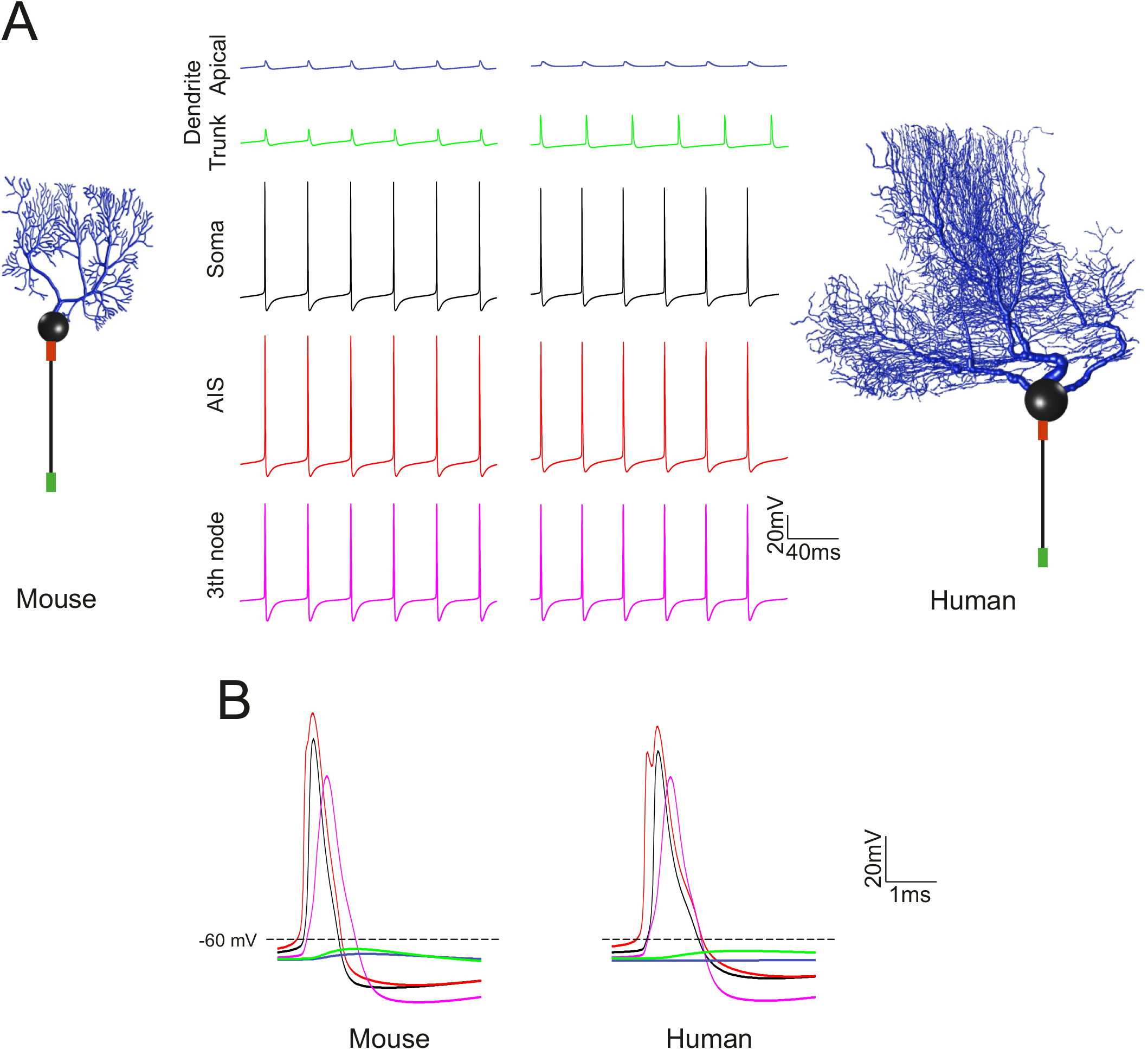
Spike propagation in PC models. (A) In all PC models, spikes are generated in the AIS and then propagate actively in the axon and passively in the dendrite, where they undergo a marked decrement and slowdown. Traces show the spontaneous spike discharge in different model compartments. The corresponding PC model morphologies are shown on the side. (B) Spikes taken from the same simulations as in A are overlayed for AIS, soma, axon, aspiny and spiny dendrites. Note that the AIS spike precedes those in any other compartment (same colors as in A).

### Responses of PC models to random synaptic inputs: burst/pause responses and the effect of synaptic inhibition

While there is abundant literature about synaptic activation in rodent cerebellar neurons, no experimental data are currently available for human neurons including PCs. In order to evaluate PC synaptic responsiveness, we made a third assumption, i.e., that the fundamental mechanisms and properties of synaptic transmission to PCs are maintained across species as suggested by comparative recordings from human pyramidal neurons ^7^. Along with this, the bursts/pause response that is observed from mice to monkeys ^19^ should also be observable in human PCs.

A burst/pause response was indeed elicited using combinations of excitatory and inhibitory synapses. An effective synaptic set able to elicit a robust response in all morphologies ^34^ was composed by 50 pf and 25 SC synapses activated by 5 impulses at 100Hz, as exemplified for a mouse and a human PC model in Fig. 5A. With this combination of excitatory and inhibitory synapses, the spike frequency increased from basal level by ~100% (n=19) in mouse and by ~60% (n=6) in human PC models. The burst frequency in human PC models increased near to the level of mice PC models by increasing pf synapses from 50 to 200 (n=6) (Fig. 5B). The intensity of burst and the length of the pause increased when inhibitory synapses were switched off. Therefore, the human PC models generated burst/pause responses like those of the mouse PC model but required 4-times more active pf to compensate for lower input resistance.

**Figure 5.**
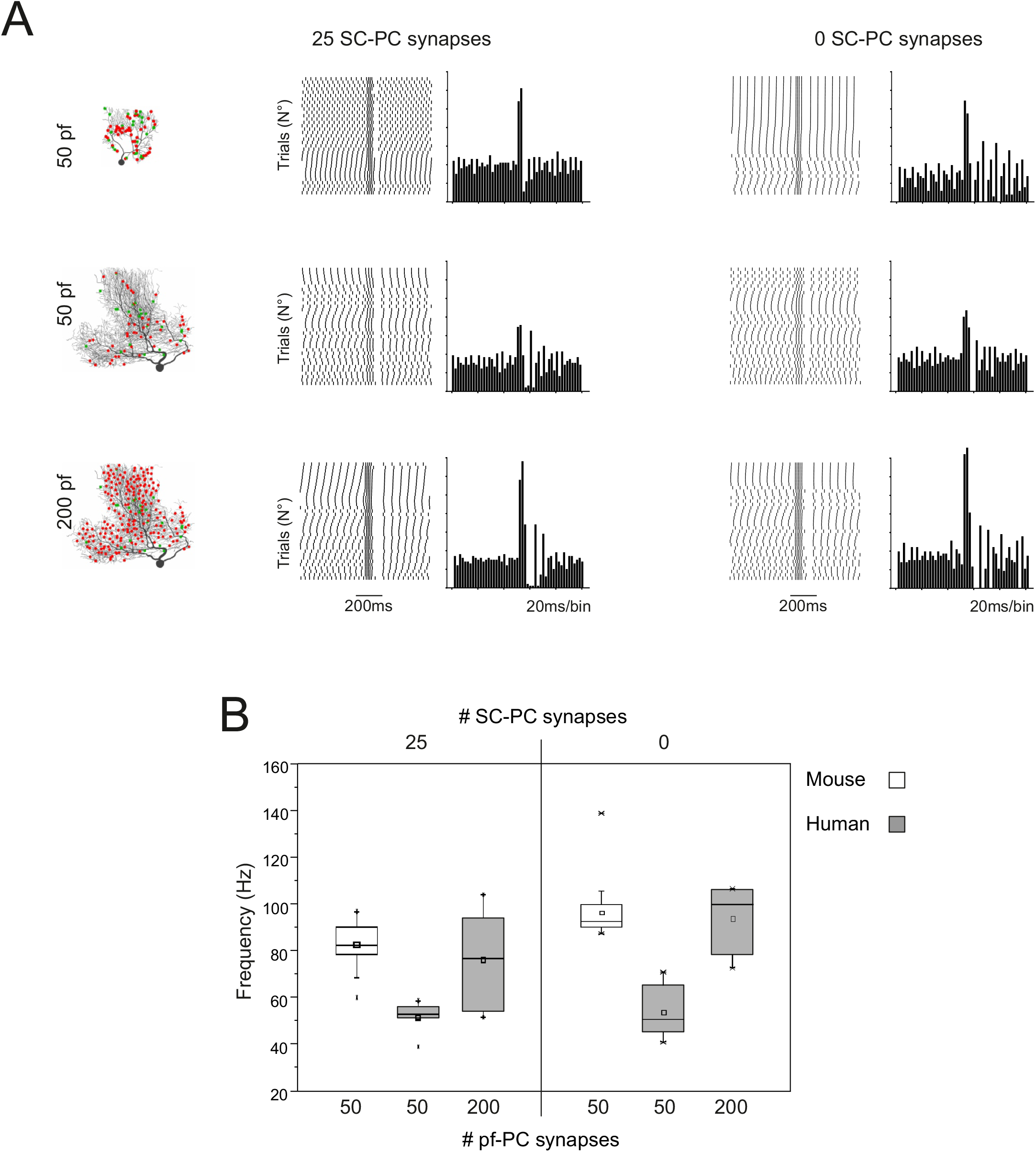
Burst/pause responses of PC models to random synaptic inputs: the effect of synaptic inhibition. (A) In a mouse and a human PC model (same as in Fig. 4), 50 pf (red dots) and 25 SC synapses (green dots) are distributed randomly on the dendritic tree ^28^. The synapses are stimulated with bursts of 5 impulses at 100Hz. Both models generate a burst/pause response that is evident in the raster plots and PSTHs. Note that the pause is more marked when inhibition is active. (B) Box plots of burst frequency using the same activation patterns as in A in all the mouse and human PC models. Note that increasing the number of pf synapses by 4 times brought the human PC model to a level of synaptic responsiveness similar to that of the mouse PC model. Note also that synaptic inhibition reduces spike frequency during the burst.

### Responses of PC models to localized synaptic inputs: dendritic independence

To assess the impact of localized synaptic inputs, the dendritic tree of PC models was stimulated on specific branches with the same synaptic activity pattern defined above (50 pf and 25SC, 5 impulses at 100Hz) (Fig. 6A). Activation of these branches effectively regulated spike/burst patterns in the soma. The independence of dendrites was explored by stimulating specific dendritic branches. Both human and mouse PC models demonstrated a remarkable degree of segregation. The membrane voltage recorded in the dendrites increased within the stimulated area but showed just marginal modifications in the other branches (Fig. 6B; cf. ref. ^28^). Considering that human have 5.5 more dendritic branches that mouse PCs (see Fig. 2C), human PCs should have an equivalent higher capacity than mouse PCs of elaborating independent input patterns in dendrites and of integrating them in the soma.

**Figure 6.**
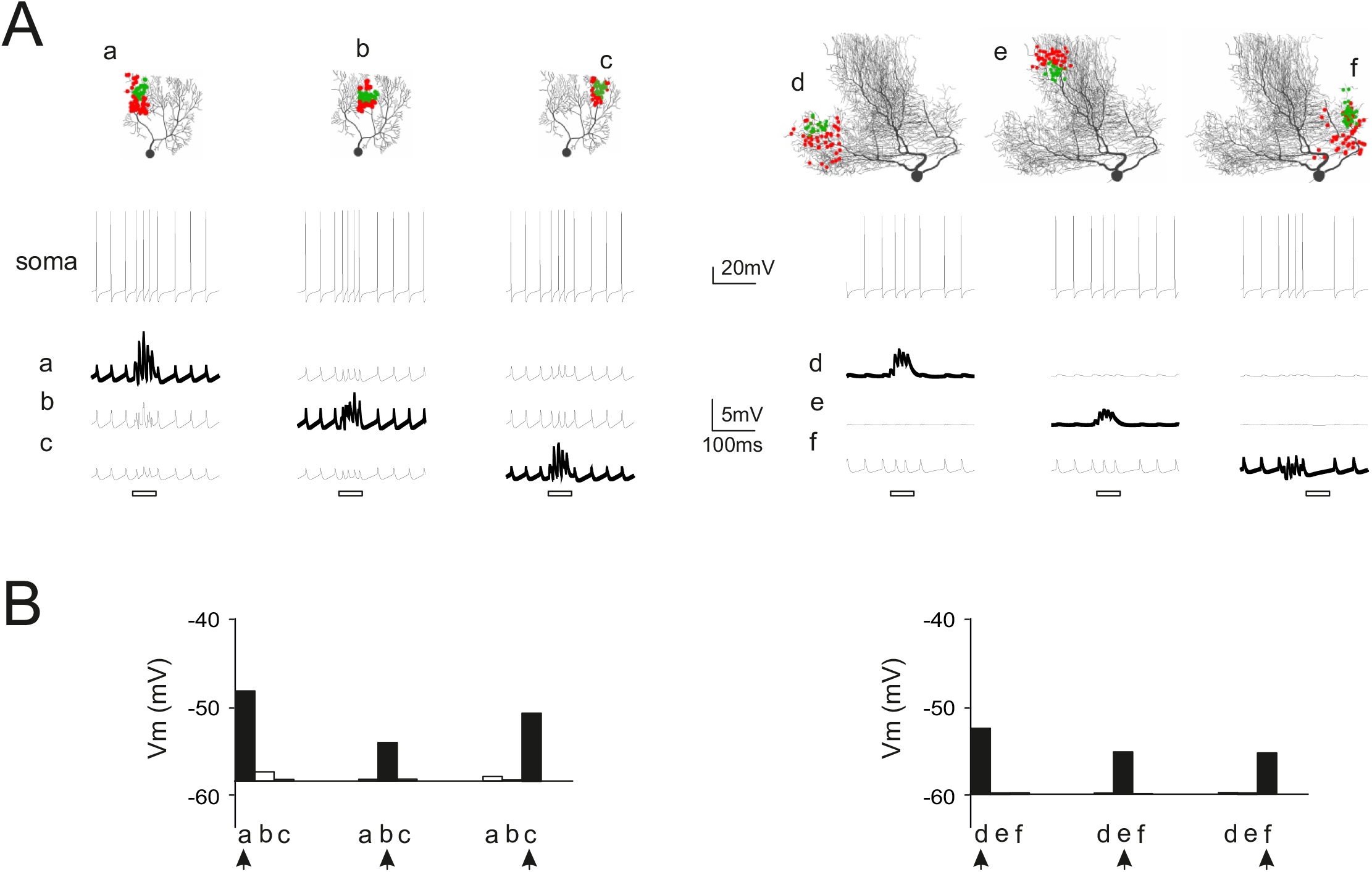
Responses of PC models to localized synaptic inputs: testing dendritic independence. (A) A mouse and a human PC model (same as in Figs 6-7) are activated by 50 pf (red dots) and 25 SC synapses (green dots) placed on different aeras of the dendritic tree (a, b, c and a1, b’, c’, respectively) ^28^. The synapses are stimulated with bursts of 5 impulses at 100Hz in each region in turn. For each stimulated region, the traces show burst/pause responses at the soma and local membrane potential changes in a compartment of the dendrites. The bars under the traces indicate the stimulation burst. (B) The histograms show membrane potential in a compartment inside each dendritic region averaged over burst duration. Note that membrane potential remarkably increases in the stimulated region (arrow) but remains almost unchanged with respect to basal level (dotted line) in the others.

### Reponses of PC models with dendritic spines

Spines may affect the local electrical properties of dendrites ^88^. Therefore, an advanced version of the PC models was developed with dendritic spines that were placed according to the density and distribution revealed by morphological analysis (see Fig. 2C) and endowed with active mechanisms according to literature (Fig. 7A). Despite the much higher computational weight (the spines increase neuronal compartments by ~100 times), models with spines were more realistic in terms of local postsynaptic mechanisms and were used to further investigate the ability of dendrites in segregating multiple synaptic input channels (Fig. 7B).

**Figure 7.**
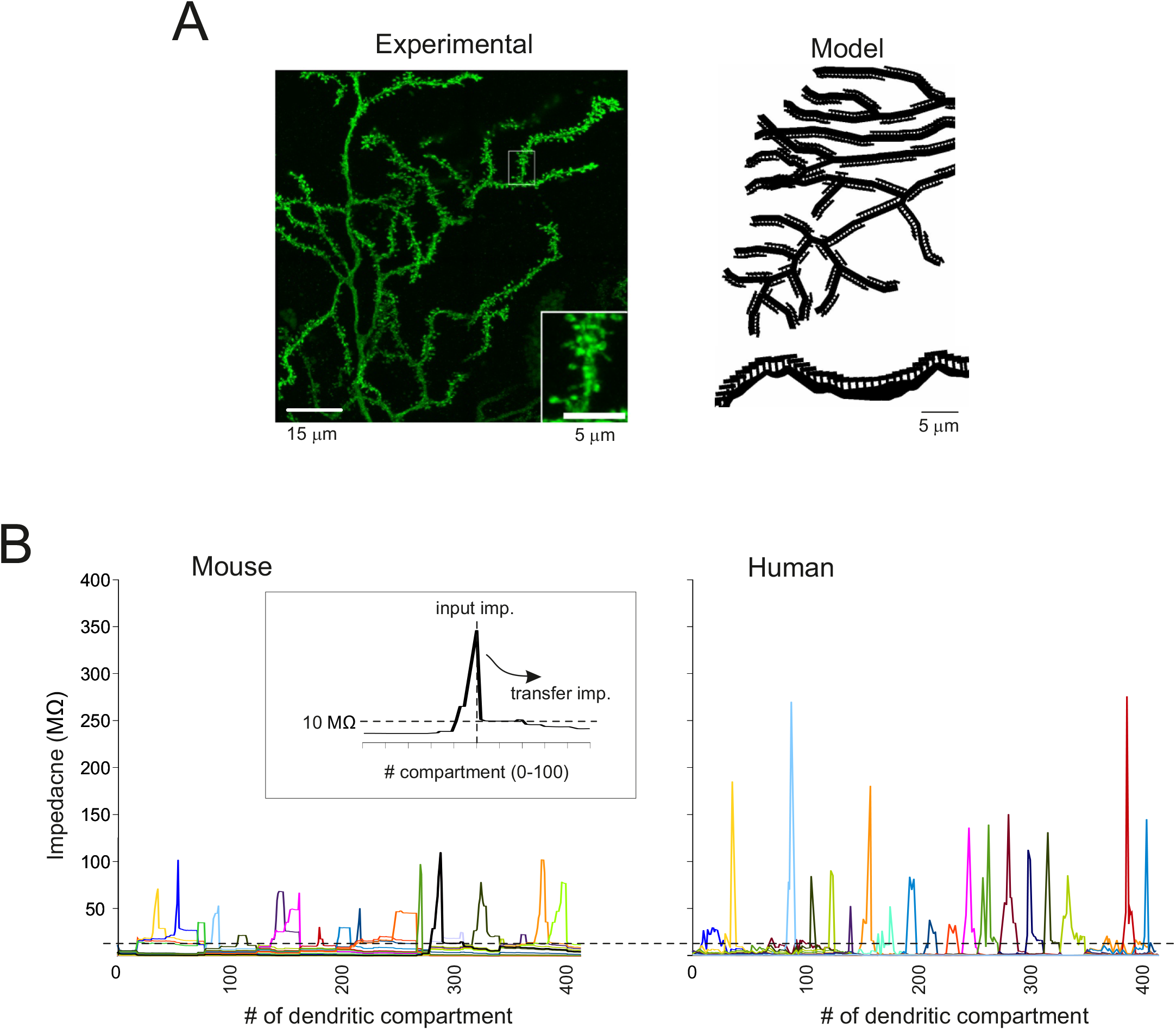
PC models with dendritic spines: re-testing dendritic independence. (A) The left panel shows a portion of the dendritic tree in a human PC intracellularly injected with Lucifer yellow. The inset shows spines at high magnification. The right panel shows the spines in the model drawn with NEURON graphics. The spine density is the same as in the experimental measurement but spines are distributed in 2D rather than 3D. (B) Examples of impedance traces calculated in a mouse and a human PC model. A 10 Hz sinusoidal current was injected into some dendrites and the recorded voltage was used to calculate the input and transfer impedance (see inset and methods). Two compartments were considered independent when the transfer impedance was below the threshold level of 10 MΩ ^84^. Note that the threshold is crossed within 10-20 compartments (e.g., see inset) and that transfer impedance in neighboring dendrites remains above threshold for longer distances in the mouse than human PC model. Note also that input impedance is higher in the human than mouse PC model reflecting the greater length of the dendrites.

A10 Hz sinusoidal current was injected in single dendrites and the membrane potential change in the same and neighboring dendrites was used to calculate input and transfer impedance (see Methods and Fig. 7B). It should be noted that input impedance was higher in human than mouse PC models, reflecting the longer pathway that currents have to travel beside the injection site.Both in human and mouse PC models, transfer impedance decreased sharply beside the current injection site. According to ^84^, when transfer impedance was below a 10 MΩ threshold, compartments were considered independent. The operation was repeated at different dendritic locations allowing to calculate an average value for the number of co-stimulated dendrites, which were 31.2±22.2 (n= 37) in mouse and 14.7±8.5 (n= 179) in human PC models. Given the total number of spiny dendrites (~250 in mice and ~750 in human PCs; Fig. 2C), the number of independent compartments turned out to be 8.0 in mouse and 51.1 in human PC models. This yields a 6.5 times higher number of independent computational elements in human than mouse PC models.

The synaptic responsiveness of PC models was assessed by delivering an input burst (5 impulses at 100 Hz) to an increasing number of pf synapses confined within the same dendrite (Fig. 8A). Local EPSPs were slower in human compared to mice PC models (Fig. 8B) probably due to the higher resistive and capacitive load that increases membrane time constant. The EPSP amplitude increased non-linearly, with comparatively smaller activation using just a few synapses followed by a rapid grow toward a plateau with more than ~50 synapses. The crosstalk between the site of origin of the synaptic response and the AIS was evaluated by measuring the changes in spike frequency at the soma (Fig. 8C). Activation of about 5 spines was enough to determine a sizeable change in instantaneous spike frequency that progressively increased with the number of active spines. With >=50 spines, the burst/pause complex appeared. Given the average number of 67 synapses/branch in human PCs and of 50 synapses/branch in mouse PCs, in both species the burst/pause response could be generated by the full activation of a single terminal branch.

**Figure 8.**
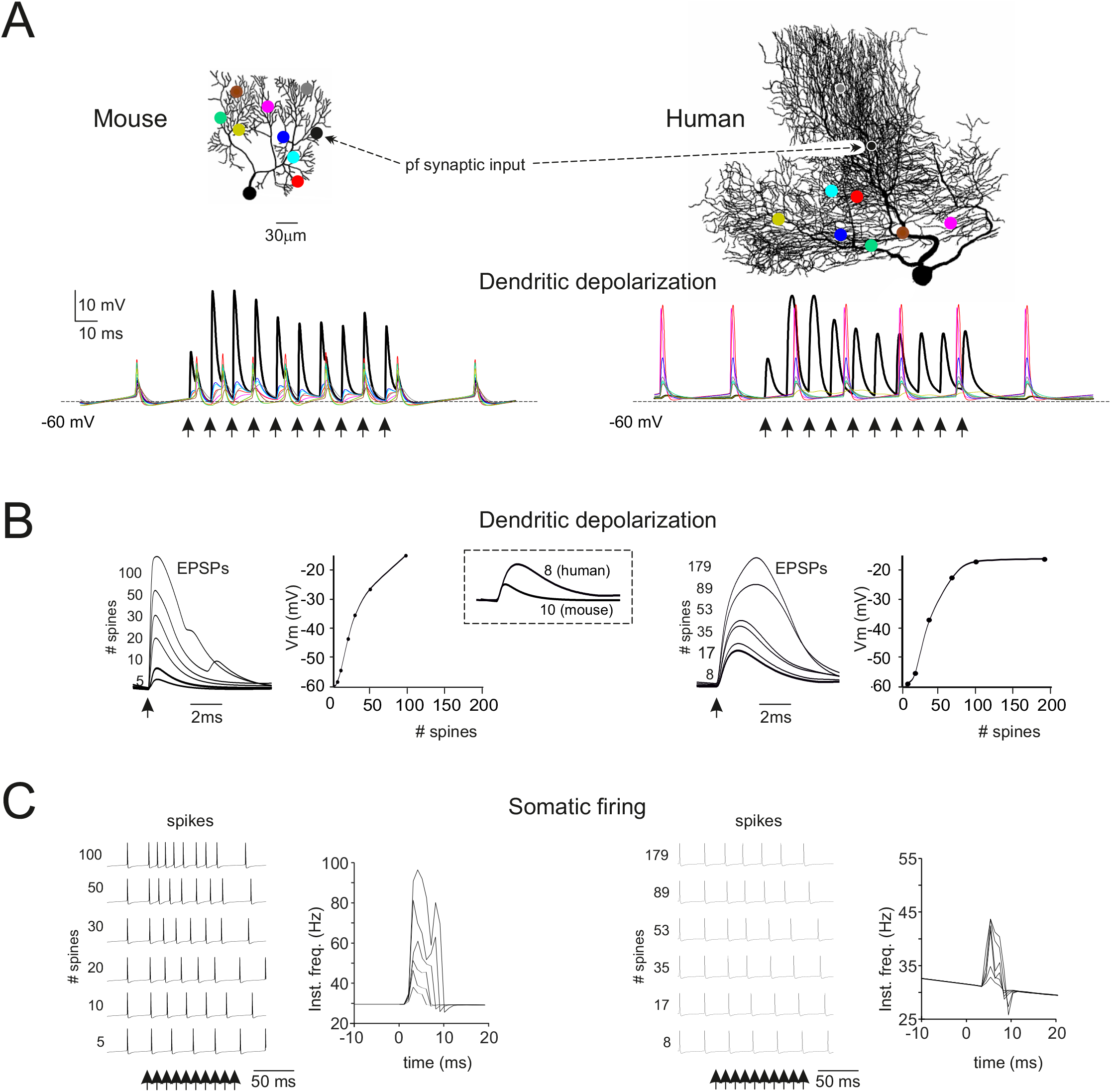
Synaptic excitation in spiny dendrites. A) A mouse and a human PC model (the same as in Figs 6-8) endowed with spines are activated by pf synapses (black dots) using bursts of 10 spikes at 100 Hz (arrows) on 50 and 89 spines, respectively The traces show responses at the stimulation site (black) and at each other site (color coded). Note short-term facilitation in pf-PC EPSPs, a much larger response at the stimulated site than in any other sites, and synchronous back-propagated spikelets visible at all sites. (B) Dendritic EPSPs generated by activating an increasing number of pf synapses (second response in a train using the same impulse pattern used in A). The traces on the left show EPSPs generated by an increasing number of spines and the plots on the right show the corresponding EPSP amplitude (the arrow indicates the stimulus). Note the sigmoidal shape of the activation curve (no response with 1 spine, rapid growth with intermediate number of spines, tendency to plateau at more than 100 spines) both in mouse and human PC models. The inset highlights the larger size and slower time course of human than mouse EPSPs. (C) Somatic spikes generated by activating an increasing number of pf synapses (same simulations as in B). The traces on the left are spike trains (the arrows indicate the stimuli) elicited by an increasing number of activated spines. Note that the number of spikes elicited by the stimuli increases with the number of active synapses, as shown in the instantaneous frequency plots on the right. Note that in A-C the profile of EPSPs and spike responses is similar in mouse and human PC models but the latter requires a larger number of active synapses.

## DISCUSSION

This paper addresses for the first time the functional and computational properties of human cerebellar PCs. PC structure and electroresponsiveness are remarkably similar to those of mouse PCs. However, human PCs have larger dendrites and a higher dendritic complexity enabling them to process more parallel fiber (pf) inputs than mouse PCs. Thus, human PCs have gained the ability of correlating a larger variety of inputs than rodent PCs extending their computational capacity into a higher dimensional space.

### Morphological and functional comparison between human and mouse PCs

The morphological growth of PCs in humans does not affect the length of terminal branches or spine density but rather it changes the number of dendritic trunks and ramification needed to scaffold and demultiplex the terminal branches. This makes neuron geometry scale-invariant (or fractal) but at the same time increases dendritic complexity (see eq. 2) and synaptic inputs. Interestingly, the presence of multiple dendritic trunks bears implications for climbing fiber connectivity and long-term synaptic plasticity ^89^. Thus, computationally, human PCs are more than just a rescaled version of the mouse or guinea pig PC ^30^, as also suggested to happen for CA1 pyramidal neurons ^90^.

The intrinsic electroresponsiveness in human and mouse PCs was similar, with regular spike firing in the depolarizing direction, sagging inward rectification in the hyperpolarizing direction, and input-output functions (the I-F relationship) increasing sub-linearly in the 100 Hz range ^11,15,17,19^. Computational simulations show that the I-F slope scales with input resistance by a factor of 2, reflecting the change in PC size. Then, both in human and mouse PCs, the spikes are generated in the AIS and backpropagate through the dendritic tree, where they are filtered down to spikelets of a few millivolts, as observed experimentally in rodents ^12,13,15^.

Simulations of dendritic activation suggest that fundamental dendritic computational properties are similar in human and mouse PCs. Low release probability causes short-term facilitation ^91^, so that sizeable dendritic responses occur when 2-3 parallel fiber spikes arrive in short sequence. Moreover, the local response grows sigmoidally with the number of spines, due to voltage-dependent calcium channel activation (cf. ^28^). In this way, coactivation of 3-5 neighboring spines is needed to generate EPSPs sufficiently large to cause a detectable acceleration of AIS spike discharge. The combination of local non-linear amplification with short-term facilitation can effectively limit transmission of sparse pf impulses, suggesting that PC dendrites operate as high-pass filters increasing signal-to-noise ratio. When a sufficient number of synapses is activated by repetitive parallel fiber inputs, a typical burst-pause response emerges as documented from rodents to monkeys ^19^. As shown in a guinea pig PC model, also in the human and mouse PC models the burst-pause responses is generated by intrinsic membrane properties ^28,36^ and is reinforced by inhibitory synaptic inputs.

### Higher computational capacity in human than mouse PCs

Two functional estimates indicate a higher dendritic complexity in human than mouse PCs: the response to synaptic input currents yields the ratio human/mouse = 5; transfer impedance measured by injecting sinusoidal 10 Hz currents yields the ratio human/mouse = 6.5. Interestingly, these two functional estimates converge toward the spine head ratio (human/mouse=5), the dendritic surface ratio (human/mouse=5.5), and the dendritic complexity ratio (human/mouse=6.5). This suggests that the increased number of contacts is almost entirely transformed into effective combinations of independent input patterns that can regulate spike generation in the soma, akin with linear encoding ^26^ in a perceptron ^25^.

The maximum computational capacity can be calculated considering the number of alternative states set up by the dendrites. The binary combinations are *K=2^n^,* where *n*=51 and *n*=8 are the numbers of independent computational elements identified in human and mouse Purkinje cells, respectively. This leads to the Shannon information *I*=Log_2_*K* of 256 bit for the mouse and 2.2*10^15^ bit for the human PCs. Due to redundancy of dendritic combinations, some output spike patterns may be mutually indistinguishable on the temporal resolution scale of the neuron and *I* estimated from Shannon’s information entropy should be considered as an upper limit. Ad-hoc simulations may allow to calculate the mutual information transfer in human and mouse PCs (e.g. see ^39–41^.

### Considerations on human PC simulations

Rodent PC models have a long tradition ^25,27–29,31–36^. In line with this, the present mouse PC models have been heavily constrained against a rich electrophysiological dataset and a wealth of literature data that were used for construction and validation. Conversely, the human PC models was uniquely based on 3D morphologies and somatic electrophysiological current-clamp recordings, which contain information about the ionic conductances determining intrinsic electroresponsiveness ^46^. The expression of equivalent channel genes in PCs across species ^87^ and the conservation of morphological features explain why the same set of ionic channels (with minor differences) could generate similar intrinsic electroresponsiveness to somatic current injection. However, some specific dendritic properties might differ. For example, pyramidal neurons have been reported to express shorter membrane time constant ^4^ and enhanced EPSP amplification in humans than mice to minimize the impact of their huge dendritic tree ^7,51^. Moreover, EPSPs generated by dendritic NMDA channels ^51^ and HVA channels ^92^ are larger in human pyramidal neurons, where they are supposed to counteract dendritic filtering. And synaptic currents in human pyramidal neurons are reported to have higher conductance for optimal synaptic transmission and integration ^51^. In another report, enhanced electrical compartmentalization of human pyramidal neuron dendrites reduced their capacity of exciting the soma ^5^. It would be interesting to see whether some sort adaptation also occur in PC dendrites, in which the high resistive and capacitive load slow down the EPSPs (e.g. see Fig. 8B) at odd with the requirements for fast processing on the millisecond scale of the cerebellar network ^93^.

### Conclusions

In conclusion, the main structural and electroresponsive properties of PCs are conserved over almost 100 million years of evolution, from rodents to humans ^54^. However, human PCs show increased dendritic complexity, which brings about a larger number of dendritic units that can regulate the neuronal output. This, in turn, causes a combinatorial explosion increasing the neuron information entropy up to 10 billion times. The higher number of independent computations that can be carried out by human than mouse PCs can be seen as the correlate of the extended cerebellar connectivity. This, in humans, involves not just sensorimotor areas but also associative areas, which engage more than 80% of all the cerebello-cortical and cortico-cerebellar fiber tracts determined by MRI tractography ^94^. The human PC would thus operate as a multimodal perceptron combining a broad information set pertaining to the sensorimotor, cognitive and emotional domains ^95^ harmonizing the cerebellar response and integrating different aspect of behavior in a high-dimensional space. It remains to be assessed whether specific adaptations of voltage-gated ionic channels, synaptic transmission, and passive membrane properties, which have been shown to improve the performance of human pyramidal neurons, also occur in human PCs.

