## Supplemental Tables and Figures for "Human outperform mouse Purkinje cells in dendritic complexity and computational capacity"

### SUPPLEMENTARY

#### Table 1 Supplementary. Morphometric parameters of mouse and human PC used for modelling.

| PC # | Dist dend | Prox_dend | Sodium dend | Tot dend | Tot dend len | Average len | Branchpoints | Sect/branch | Branch order | Branch angle |
|---|---|---|---|---|---|---|---|---|---|---|
| Mouse 1 | 197 | 20 | 1 | 218 | 2685.3 | 12.3 | 109 | 2 | 16 | 1.2 |
| Mouse 2 | 319 | 16 | 2 | 337 | 3547.4 | 10.5 | 171 | 1.8 | 18 | 1.2 |
| Mouse 3 | 214 | 16 | 1 | 231 | 2677.2 | 11.6 | 115 | 2.0 | 19 | 1.1 |
| Mouse 4 | 218 | 17 | 3 | 238 | 2469.8 | 10.4 | 119 | 2 | 19 | 1.3 |
| Mouse 5 | 320 | 17 | 2 | 339 | 3178.8 | 9.4 | 172 | 2.0 | 20 | 1.3 |
| Mouse 6 | 258 | 11 | 1 | 270 | 2805.4 | 10.4 | 135 | 2 | 17 | 1.2 |
| Mouse 7 | 288 | 19 | 6 | 313 | 3054.9 | 9.8 | 157 | 2.0 | 20 | 1.3 |
| Mouse 8 | 172 | 11 | 2 | 185 | 2301.4 | 12.4 | 94 | 2.0 | 16 | 1.2 |
| Mouse 9 | 178 | 11 | 2 | 191 | 2236.4 | 11.7 | 98 | 2.0 | 16 | 1.2 |
| Mouse 10 | 433 | 26 | 1 | 460 | 4464.3 | 9.7 | 229 | 2.0 | 25 | 1.2 |
| Mouse 11 | 82 | 8 | 2 | 92 | 1340.5 | 14.6 | 47 | 2.0 | 12 | 1.0 |
| Mouse 12 | 310 | 24 | 4 | 338 | 3349.5 | 9.9 | 171 | 2.0 | 19 | 1.2 |
| Mouse 13 | 124 | 14 | 1 | 139 | 1826.4 | 13.1 | 71 | 2.0 | 15 | 1.2 |
| Mouse 14 | 228 | 14 | 1 | 243 | 2584.4 | 10.6 | 124 | 2.0 | 20 | 1.2 |
| Mouse 15 | 261 | 15 | 3 | 279 | 2773.9 | 9.9 | 140 | 2.0 | 20 | 1.2 |
| Mouse 16 | 257 | 22 | 1 | 280 | 2800.0 | 10.0 | 141 | 2.0 | 22 | 1.2 |
| Mouse 17 | 341 | 17 | 2 | 360 | 3671.3 | 10.2 | 183 | 2.0 | 17 | 1.2 |
| Mouse 18 | 223 | 14 | 5 | 242 | 2470.8 | 10.2 | 124 | 2.0 | 20 | 1.2 |
| Mouse 19 | 218 | 15 | 2 | 235 | 2631.4 | 11.2 | 118 | 2.0 | 18 | 1.1 |
| Human 1 | 1678 | 93 | 35 | 1806 | 51426.7 | 28.5 | 925 | 2.0 | 42 | 1.3 |
| Human 2 | 354 | 33 | 6 | 393 | 9817.3 | 25.0 | 199 | 2.0 | 28 | 1.2 |
| Human 3 | 303 | 30 | 2 | 335 | 9019.7 | 27.0 | 172 | 2.0 | 18 | 1.2 |
| Human 4 | 285 | 35 | 7 | 327 | 8680.9 | 27.0 | 164 | 2.0 | 18 | 1.2 |
| Human 5 | 526 | 26 | 8 | 560 | 15873.3 | 28.3 | 294 | 2.0 | 22 | 1.3 |
| Human 6 | 1145 | 40 | 52 | 1237 | 26183.8 | 21.2 | 642 | 2.0 | 39 | 1.2 |

Table 1. Morphometric parameters obtained from NEURON (columns 1-6) and Tree toolbox (columns 7-10) in each one of the reconstructed neuronal morphologies used for modelling.
Dist dend: Number of spiny terminal dendrites
Prox dend: Number of aspiny proximal dendrites
Sodium dend: Number of aspiny proximal dendrites endowed with sodium channels
Tot dend: Total number of dendritic sections
Tot dend len: Total dendritic length in µm
Average len: Average length of dendritic sections
Branch points: Total number of branch points
Sect/branch: Number of sections for each branch point
Branch order: Order of the branches obtained using the tree toolbox

Branch angle: Average angle (in degrees) of the section departing from a branch point

**Table 2 Supplementary. Optimisation results of mouse and human PC models.**

| Morphology | Individuals | Valid I/O | % I/O | Valid AIS | % AIS | Valid Synaptic activity | % Syn |
|---|---|---|---|---|---|---|---|
| Mouse 1 | 576 | 491 | 85.24 | 491 | 85.24 | 383 | 66.49 |
| Mouse 2 | 576 | 531 | 92.18 | 531 | 92.18 | 89 | 15.45 |
| Mouse 3 | 576 | 384 | 66.66 | 384 | 66.66 | 168 | 29.16 |
| Mouse 4 | 576 | 485 | 84.20 | 485 | 84.20 | 128 | 22.22 |
| Mouse 5 | 576 | 567 | 98.43 | 565 | 98.09 | 394 | 68.40 |
| Mouse 6 | 576 | 522 | 90.62 | 522 | 90.62 | 457 | 79.34 |
| Mouse 7 | 576 | 428 | 74.30 | 428 | 74.30 | 243 | 42.18 |
| Mouse 8 | 576 | 340 | 59.02 | 340 | 59.02 | 107 | 18.57 |
| Mouse 9 | 576 | 569 | 98.78 | 567 | 98.43 | 17 | 2.951 |
| Mouse 10 | 576 | 221 | 38.36 | 221 | 38.36 | 68 | 11.80 |
| Mouse 11 | 576 | 538 | 93.40 | 233 | 40.45 | 210 | 36.45 |
| Mouse 12 | 576 | 542 | 94.09 | 542 | 94.09 | 7 | 1.215 |
| Mouse 13 | 576 | 354 | 61.45 | 354 | 61.45 | 60 | 10.41 |
| Mouse 14 | 576 | 394 | 68.40 | 394 | 68.40 | 124 | 21.52 |
| Mouse 15 | 576 | 393 | 68.22 | 393 | 68.22 | 68 | 11.80 |
| Mouse 16 | 576 | 565 | 98.09 | 565 | 98.09 | 350 | 60.76 |
| Mouse 17 | 576 | 114 | 19.79 | 114 | 19.79 | 41 | 7.118 |
| Mouse 18 | 576 | 455 | 78.99 | 455 | 78.99 | 247 | 42.88 |
| Mouse 19 | 576 | 519 | 90.10 | 519 | 90.10 | 380 | 65.97 |
| Total | 10944 | 8412 | 76.86 | 8103 | 74.04 | 3541 | 32.35 |
| Human 1 | 576 | 509 | 88.36 | 509 | 88.36 | 246 | 42.70 |
| Human 2 | 576 | 404 | 70.13 | 145 | 25.17 | 70 | 12.15 |
| Human 3 | 576 | 499 | 86.63 | 482 | 83.68 | 458 | 79.51 |
| Human 4 | 576 | 575 | 99.82 | 217 | 37.67 | 22 | 3.81 |
| Human 5 | 576 | 559 | 97.04 | 552 | 95.83 | 128 | 22.22 |
| Human 6 | 576 | 512 | 88.88 | 458 | 79.51 | 15 | 2.60 |
| Total | 19584 | 15344 | 78.34 | 14035 | 71.66 | 5967 | 30.46 |

Table 2. The table illustrates the optimization and validation process.
Individuals: total number of individuals in the last generation of optimization
Valid I/O: number of individuals with a correct Input Output relationship
% I/O: percentage of individuals valid for the Input Output relationship
Valid AIS: number of individuals valid for the absence of intrinsic activity if the sodium channels are missing from the Axon Initial Segment.
% AIS: percentage of individuals valid for the absence of intrinsic activity if the sodium channels are missing from the Axon Initial Segment.
Valid Synaptic activity: Number of individuals validated for the synaptic activity.
% Syn: Percentage of individuals validated for synaptic activity.

**Table 3 Supplementary. Ionic channels and maximum conductances ranges.**

| Ionic channel type | Location | Conductance ranges (mA/cm$^2$) |
|---|---|---|
| Kv3.3 | 0 µm<= Dendrites <= 1.6 µm | 0.009 - 0.013 |
| Kv4.3 | | 0.0008 - 0.001 |
| Cav2.1 | | 7.5e-3 - 2e-2 |
| Cav3.1 | | 4e-6 - 8e-6 |
| Cav3.2 | | 0.00085 - 0.002 |
| Cav3.3 | | 0.0001 - 0.00017 |
| Kca1.1 | | 2.5e-2 - 4e-2 |
| Kca2.2 | | 7e-4 - 1e-3 |
| HCN1 | | 0.000001 - 0.0000032 |
| Kv1.1 | Dendrites >= 1.6 µm | 0.001 - 0.0013 |
| Kv1.5 | | 0.9e-4 - 1.5e-4 |
| Kir2.3 | | 0.000008 - 0.00002 |
| Kca3.1 | | 0.0025 - 0.006 |
| Nav1.6 | Dendrites >= 3.3 µm | 0.0145, 0.016 |
| Nav1.6 | Soma | 0.21 - 0.25 |
| Kv1.1 | | 0.001 - 0.003 |
| Kv1.5 | | 3.5e-4 - 5e-4 |
| Kv3.4 | | 0.07 - 0.1 |
| Kir2.3 | | 0.000015 - 0.00007 |
| Cav2.1 | | 2.5e-4 - 5e-4 |
| Cav3.1 | | 4e-6 - 9e-6 |
| Cav3.2 | | 0.0006 - 0.001 |
| Cav3.3 | | 0.00009 - 0.00015 |
| Kca1.1 | | 0.008 - 0.015 |
| Kca2.2 | | 8e-4 - 1.5e-3 |
| Kca3.1 | | 0.007 - 0.011 |
| HCN1 | | 0.001 - 0.009 |
| Nav1.6 | AIS | 1.75 - 2 |
| Kv3.4 | | 0.008 - 0.011 |
| Cav2.1 | | 1e-4 - 4e-4 |
| Cav3.1 | | 5e-6 - 1e-5 |
| Kv1.1 | Para AIS | 0.008 - 0.02 |

| | | |
|---|---|---|
| Nav1.6 | Nodes of Ranvier | 0.025 - 0.04 |
| Kv3.4 | | 0.01 - 0.015 |
| Cav2.1 | | 1e-4 - 3e-4 |
| Cav3.1 | | 1e-5 - 2e-5 |

Ionic channel type: ionic channel type based on an international classification (Yu et al., 2005)
Location: the location along the morphology were the ionic channels were placed.
Conductance ranges (mA/cm$^2$): The parameter ranges were used as priors for the optimization process and were the same both in mouse and human PC models.

**Table 4 Supplementary. Ionic channels maximum conductance following optimization.**

| Ionic channel type | Location | Max conductance (mA/cm$^2$) |
|---|---|---|
| Kv3.3 | 0 µm<= Dendrites <= 1.6 µm | 0.01065 ± 0.0012937 |
| Kv4.3 | | 0.00102 ± 0.0001928 |
| Cav2.1 | | 0.00099 ± 0.0004428 |
| Cav3.1 | | 7.11566E-06 ± 7.32095E-07 |
| Cav3.2 | | 0.00181 ± 9.11506E-05 |
| Cav3.3 | | 0.00014 ± 3.17121E-05 |
| Kca1.1 | | 0.03915 ± 0.0085483 |
| Kca2.2 | | 0.00093 ± 0.0001712 |
| HCN1 | | 2.05563E-06 ± 6.62379E-07 |
| Kv1.1 | Dendrites >= 1.6µm | 0.00122 ± 3.1379E-05 |
| Kv1.5 | | 0.00012 ± 2.1532E-05 |
| Kir2.3 | | 1.23279E-05 ± 4.28247E-06 |
| Kca3.1 | | 0.00370 ± 0.0009820 |
| Nav1.6 | Dendrites >= 3.3 | 0.0151 ±0.0005024 |
| Nav1.6 | Soma | 0.21720 ± 0.0236269 |
| Kv1.1 | | 0.00174 ± 0.0007324 |
| Kv1.5 | | 0.00048 ± 0.0001599 |
| Kv3.4 | | 0.07377 ± 0.0054591 |
| Kir2.3 | | 3.56706E-05 ± 2.15094E-05 |
| Cav2.1 | | 0.00035 ± 8.84837E-05 |
| Cav3.1 | | 6.95137E-06 ± 1.79487E-06 |
| Cav3.2 | | 0.00090 ± 9.3038E-05 |
| Cav3.3 | | 0.00012 ± 1.93729E-05 |
| Kca1.1 | | 0.01186 ± 0.0025027 |
| Kca2.2 | | 0.00120 ± 0.0003136 |
| Kca3.1 | | 0.00861 ± 0.0015543 |
| HCN1 | | 0.00225 ± 0.0027538 |
| Nav1.6 | AIS | 1.21748 ± 0.1464244 |
| Kv3.4 | | 0.00981 ± 0.0008169 |

| | | |
|---|---|---|
| Cav2.1 | | 0.00022 ± 9.69215E-05 |
| Cav3.1 | | 6.50376E-06 ± 1.59735E-06 |
| Kv1.1 | Para AIS | 0.01334 ± 0.0039573 |
| Nav1.6 | Nodes of Ranvier | 0.02938 ± 0.0028718 |
| Kv3.4 | | 0.01185 ± 0.0012497 |
| Cav2.1 | | 0.00018 ± 6.80297E-05 |
| Cav3.1 | | 1.53709E-05 ± 3.31895E-06 |

Ionic channel type: ionic channel type based on an international classification (Yu et al., 2005)
Location: the location along the morphology were the ionic channels were placed.
Max conductance ($mA/cm^2$): Conductances ranges obtained after optimization in mouse (n=3) and human (n=3) PC models.

**Fig. 1 Supplementary - Intracellular injection and reconstruction of human PCs**

A and B show a dorsal view of the human cerebellum (AB6 case) and a tissue block (B top) obtained from the vermis region of the anterior and posterior lobes. C shows an example of the vibratome sections that were used to intracellularly inject PCs with Lucifer yellow. D, low power conventional fluorescence photomicrograph through a cerebellar folia showing examples of PCs (arrows) intracellularly injected with Lucifer yellow. E show an intracellularly injected PC (arrow), corresponding to the squared zone in D, that was subsequently scanned by confocal microscopy (shown in F as a z-projection image) and reconstructed with Neurolucida software (G). Scale bar, shown in G, indicates 16 mm in A and B, 4 mm in C, 490 µm in D, 145 µm in E, and 65 µm in F and G.

**Fig. 2 Supplementary – Counting primary dendrites on human Purkinje cells.** Top: representative examples of human cerebellar cortex tissue sections with stained PCs. Primary dendrites can generally be clearly seen when the sectioning angle matches the translobular plane. Bottom: histogram summarizing results from 4 different tissue samples; three additional tissue samples were excluded because PC morphologies were too far degraded before tissue fixation and staining did not show primary dendrites clearly. The figure shows sections of 30 µm, 40 µm and 50 µm respectively with 1:5000 primary antibody concentration. Human post mortem male brain of age 61 fixed with 4% PFA.

METHODS: Fixed human brain tissue specimens were obtained through the NIH NeuroBioBank following a request for a preliminary study on this topic. Specimens were prepared at various times post mortem and using either paraformaldehyde or formalin as fixative. Following visual inspection to determine the best possible translobular sectioning plane, a small piece was cut from each specimen and washed in PBS. A randomly chosen subset of these tissues was placed in 30% sucrose in PBS, embedded in O.C.T. compound (brand) and cooled to -80°C before being sectioned to 20 - 50 um thickness using a cryostat, while the remaining tissues were sectioned to 50 - 100 um thickness using a vibratome. Tissues were collected and stored free-floating in PBS until staining, which was performed using a primary antibody targeting the calbindin-D28K protein (locally specific to Purkinje neurons; Swant) and a matching far-red secondary antibody (donkey-anti-rabbit-AlexaFluor-647). The general procedure was as follows: first, the tissues were rinsed 4 times in PBS for 4 minutes, and then incubated in a blocking buffer containing Normal Donkey Serum, PBS and 0.3% Triton x-100. 400ul of blocking buffer for 60 minutes at room temperature. Slices were then incubated with the primary antibody (diluted 1:2000 or 1:5000 in the blocking buffer) overnight at 4C°. On the next day the sections were rinsed 4 times 4 minutes with PBS, and then incubated in blocking solution with the secondary antibody (donkey-anti-rabbit-AlexaFluor-647) added at a concentration of 1:1000 for 90 minutes at 4C° in the dark and finally rinsed in PBS 4 times for 4 minutes and stored in PBS Azide (no more than 36 hours) until mounting.

Slices were mounted on glass microscopy slides using fine brushes, taking care that folia would be lying down flat and not get twisted. Once dried thoroughly, a few drops of mounting medium (ProLong™ Diamond Antifade Mountant or ThermoScientific PermaFluor Aqueous Mounting

Medium) were placed on each slide and the slides were covered using coverslips and allowed to cure according to the mounting medium's manufacturers' recommendations (30min - 48hrs). After the curing period, the slices were imaged using an Olympus Bright Field Microscope. Areas of interest were identified under 4x magnification, and subsequently captured using a 40x magnification objective. Images were viewed and post-processed to adjust brightness and contrast using ImageJ (Schneider et al., 2012), and 8 images with particularly clear single PN layers in the right orientation were selected for further analysis; only sections in which primary dendrites could be counted for >80% of PNs were used. Altogether, 350 PNs were examined across 4 different tissue samples, and 569 primary dendrites were counted on 297 PNs. The resulting counts were logged per image-section and from these counts, the percentage of PNs with multiple primary dendrites was calculated.

**Fig. 3 Supplementary - Mouse and human morphologies.**

The figure illustrates the entire dataset of morphologies used in the construction and validation of the PCs models.

**Fig. 4 Supplementary - Ionic channel types and distribution**

The figure illustrates the distribution of ionic channels and Ca buffer in the PC models. The ionic channels mechanism were taken from a previous model (Masoli et al., 2015; Masoli and D'Angelo, 2017) and updated to reflect new experimental data. The KCa2.2 channel was distributed over the entire dendritic tree, whereas Kv1.1 was restricted to proximal dendrites and Kv1.5 to proximal dendrites and soma. Specific ionic channels were inserted in spines according to literature.

**Fig. 5 Supplementary - Optimization process**

The panel shows the progress of optimization through subsequent generations. The fitness values for the 25 PC models tend toward zero in just 7 generations showing improved matching between experimental data and modelling results.

**Fig. 6 Supplementary – Ramp current injection**

(A) The box plot shows statistics of spontaneous firing frequency for all the mouse and human PC models.

(B) I-F relationships for all PC models covered in this study. The I-F relationships shows similar shape in human and mice PCs.

(C) A ramp current injection, from 0 to 1.6 nA in mouse and from 0 to 4.8 nA in human PC models, reproduced a typical PC voltage response (Williams et al., 2002). A marked frequency reduction appears in the falling branch of the ramp, causing an asymmetric instantaneous frequency profile.

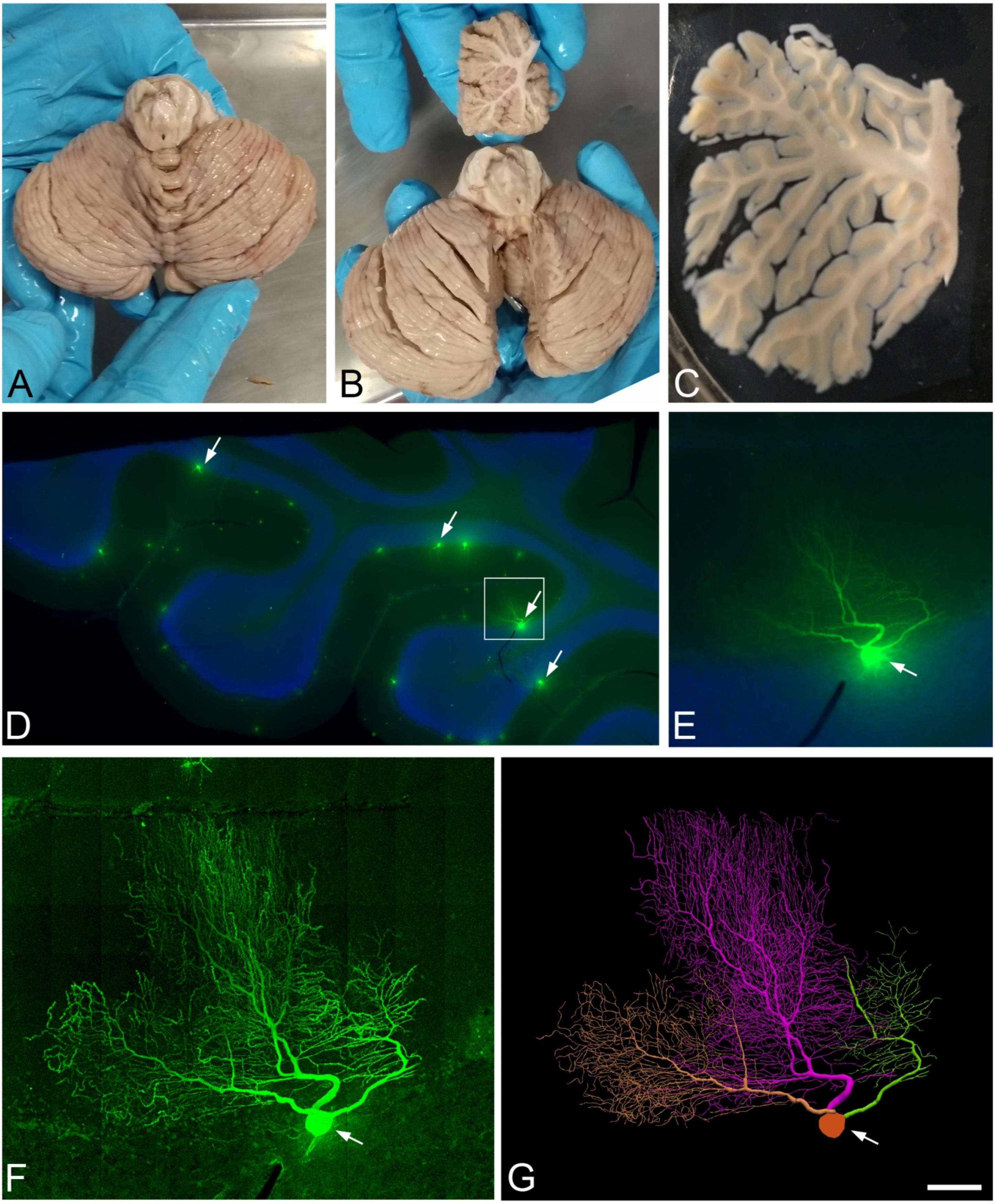
A
B
C
D
E
F
G

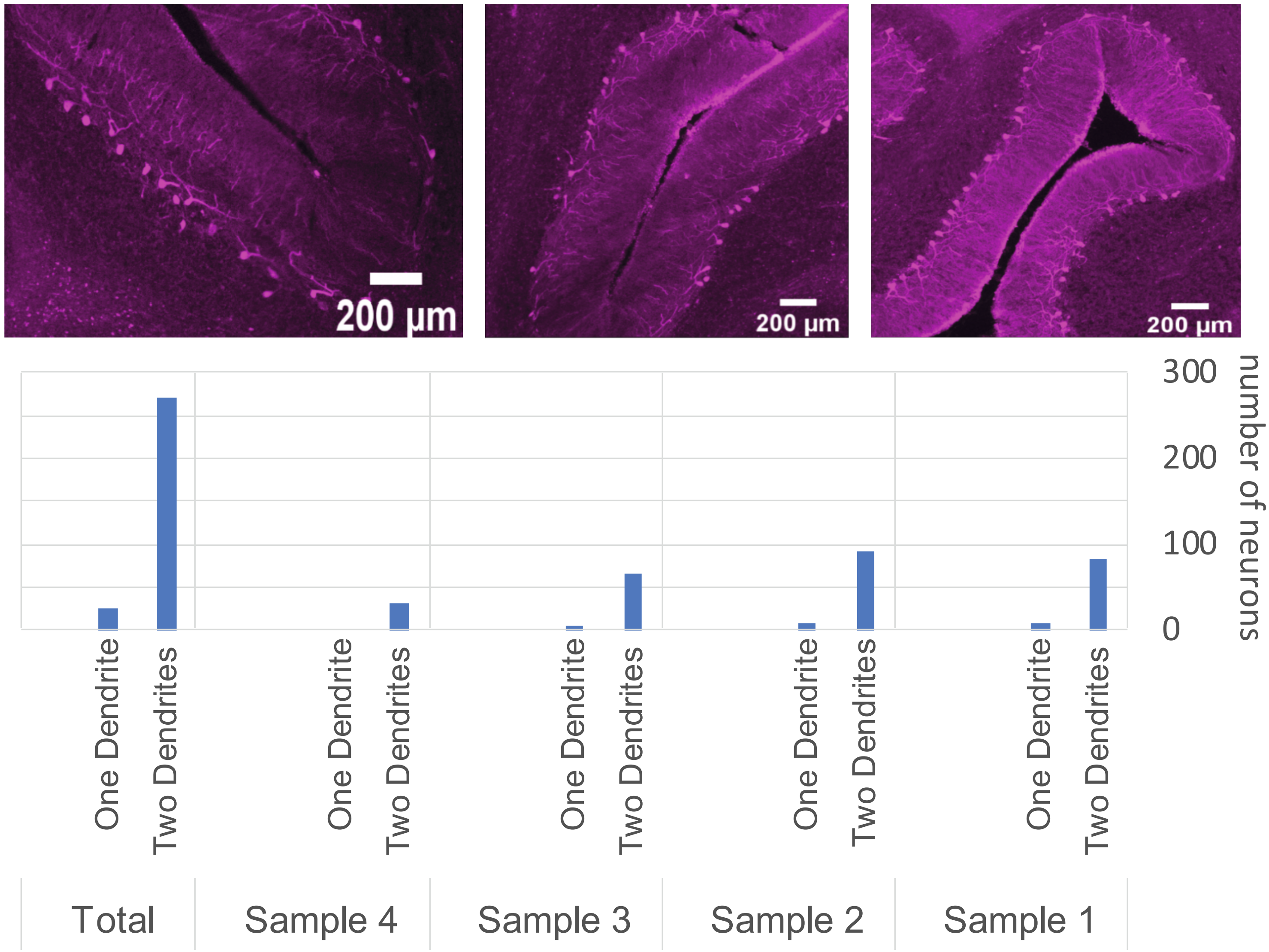

200 µm
200 µm
200 µm
300
200
100
0
number of neurons
One Dendrite
Two Dendrites
Total
Sample 4
Sample 3
Sample 2
Sample 1

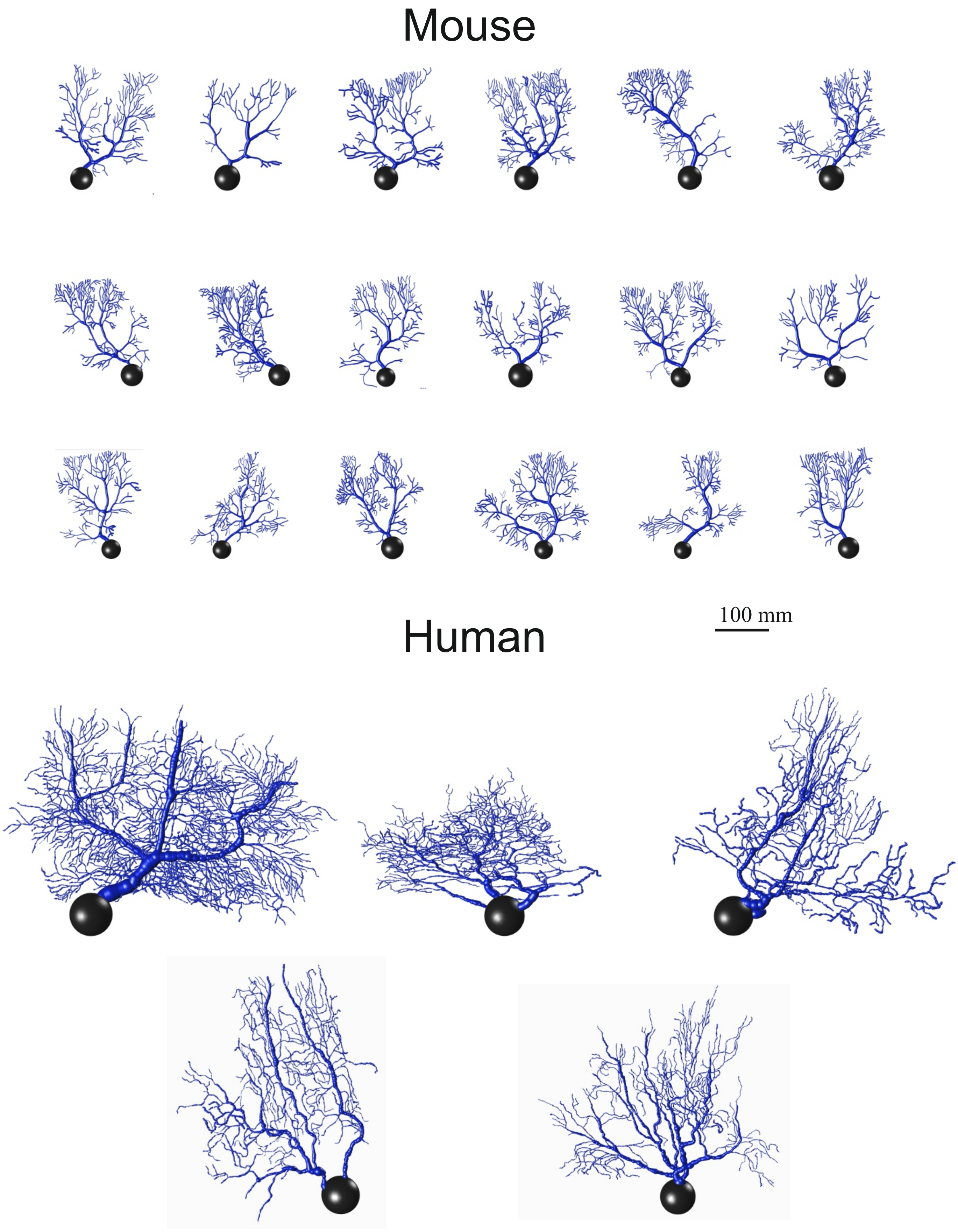
Mouse
100 mm
Human

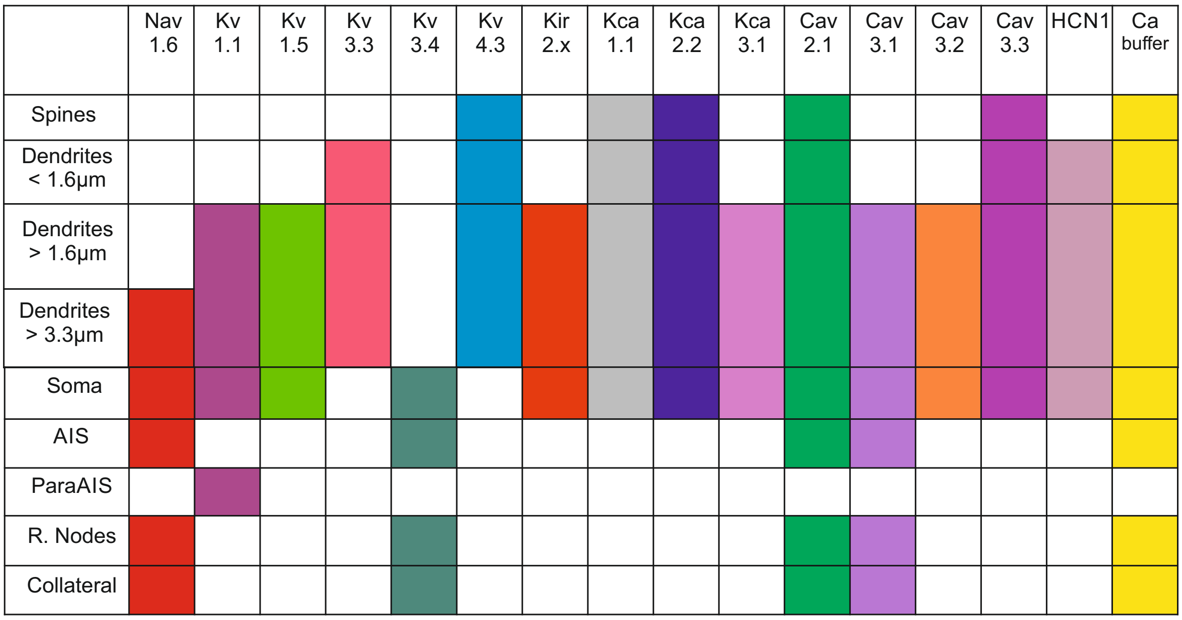

Nav 1.6
Kv 1.1
Kv 1.5
Kv 3.3
Kv 3.4
Kv 4.3
Kir 2.x
Kca 1.1
Kca 2.2
Kca 3.1
Cav 2.1
Cav 3.1
Cav 3.2
Cav 3.3
HCN1
Ca buffer
Spines
Dendrites < 1.6μm
Dendrites > 1.6μm
Dendrites > 3.3μm
Soma
AIS
ParaAIS
R. Nodes
Collateral

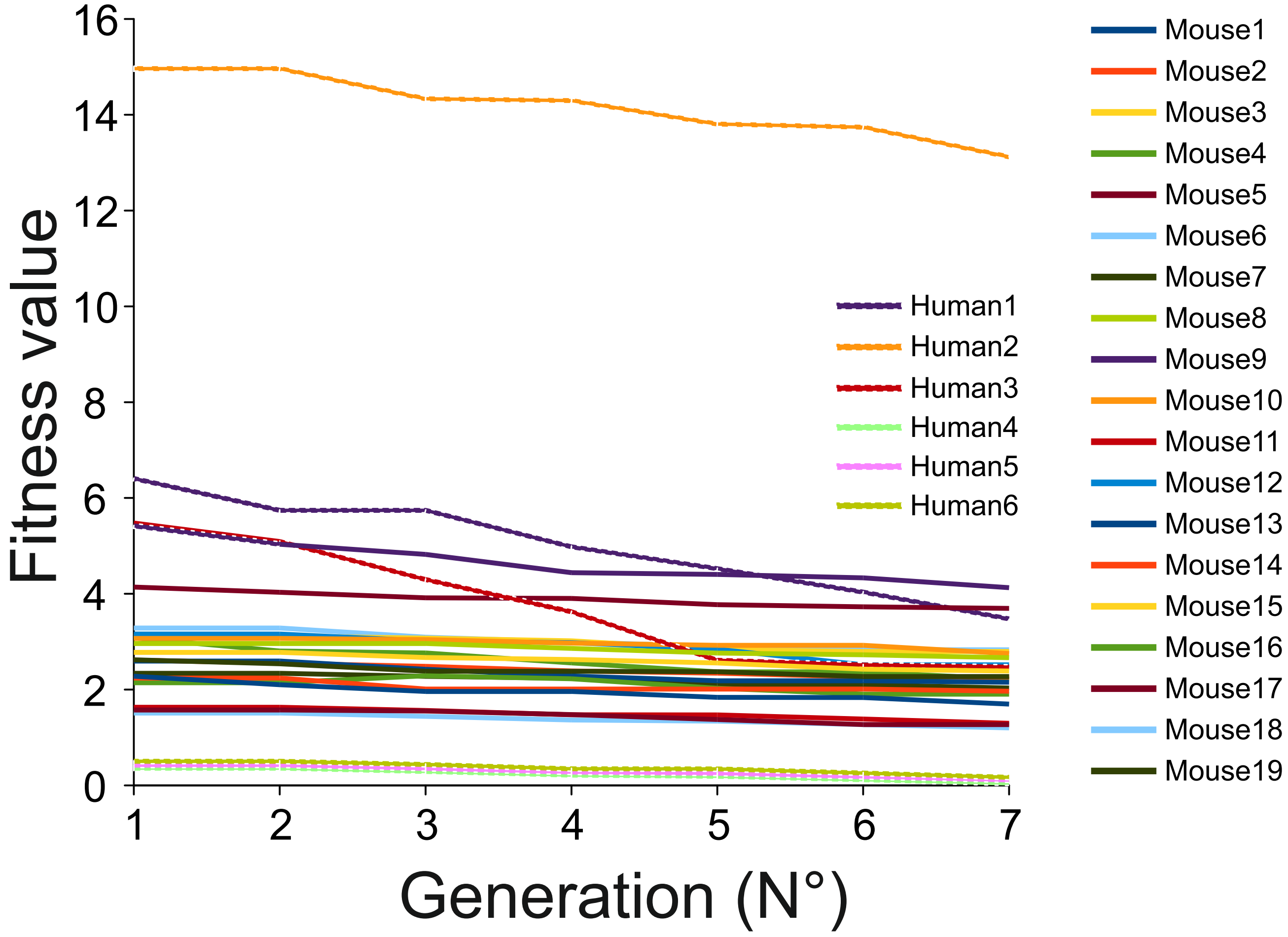

Fitness value
16
14
12
10
8
6
4
2
0
1
2
3
4
5
6
7
Generation (N°)
Human1
Human2
Human3
Human4
Human5
Human6
Mouse1
Mouse2
Mouse3
Mouse4
Mouse5
Mouse6
Mouse7
Mouse8
Mouse9
Mouse10
Mouse11
Mouse12
Mouse13
Mouse14
Mouse15
Mouse16
Mouse17
Mouse18
Mouse19

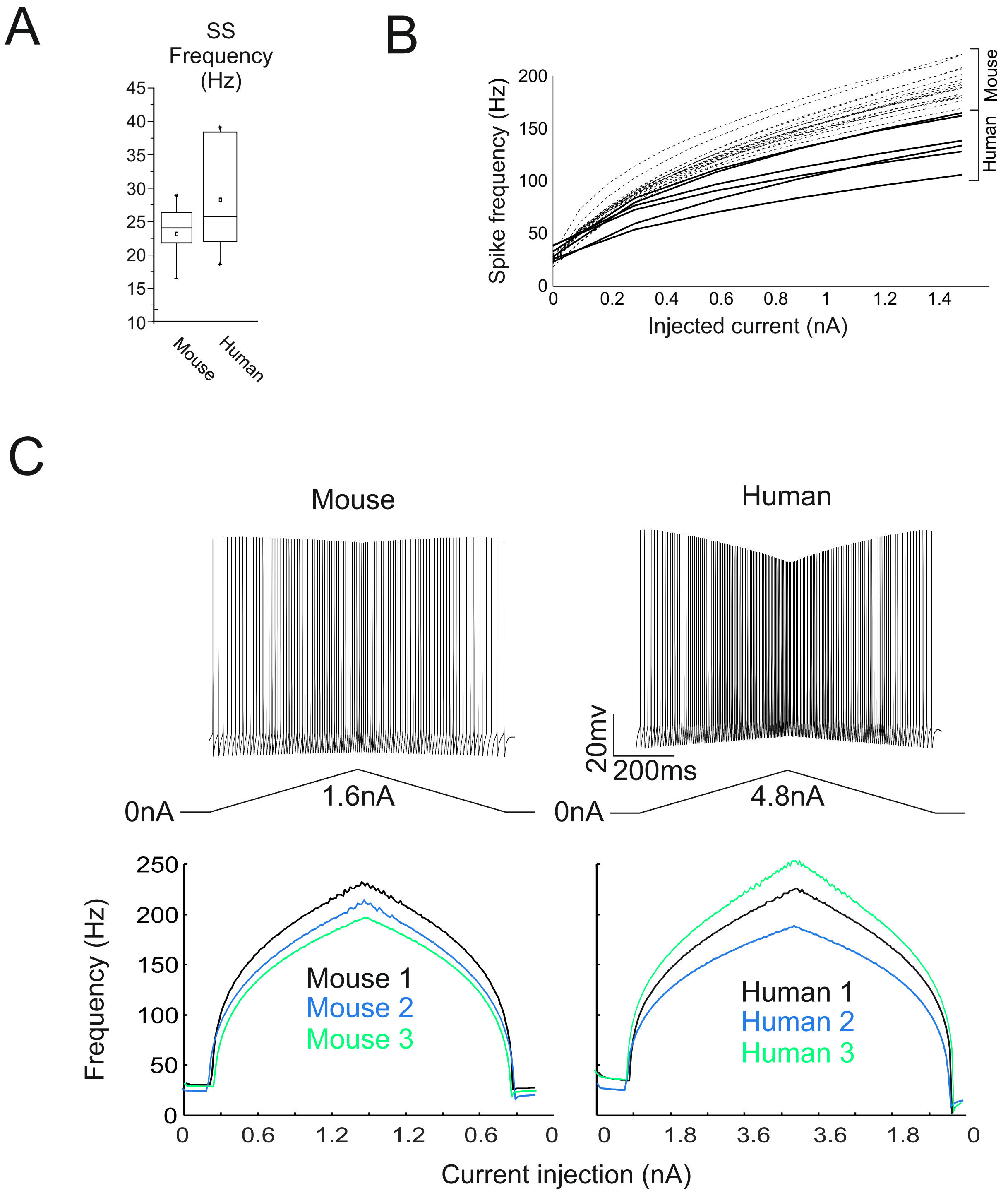

A
SS Frequency (Hz)
45
40
35
30
25
20
15
10
Mouse
Human
B
Spike frequency (Hz)
200
150
100
50
0
0
0.2
0.4
0.6
0.8
1
1.2
1.4
Injected current (nA)
Mouse
Human
C
Mouse
Human
20mv
200ms
0nA
1.6nA
0nA
4.8nA
Frequency (Hz)
250
200
150
100
50
0
Mouse 1
Mouse 2
Mouse 3
Human 1
Human 2
Human 3
0
0.6
1.2
1.2
0.6
0
0
1.8
3.6
3.6
1.8
0
Current injection (nA)
